# Genotype networks of 80 quantitative *Arabidopsis thaliana* phenotypes reveal phenotypic evolvability despite pervasive epistasis

**DOI:** 10.1101/2020.02.26.966390

**Authors:** Gabriel Schweizer, Andreas Wagner

## Abstract

We study the genotype-phenotype maps of 80 quantitative phenotypes in the model plant *Arabidopsis thaliana,* by representing the genotypes affecting each phenotype as a genotype network. In such a network, each vertex or node corresponds to an individual’s genotype at all those genomic loci that affect a given phenotype. Two vertices are connected by an edge if the associated genotypes differ in exactly one nucleotide. The 80 genotype networks we analyze are based on data from genome-wide association studies of 199 *A. thaliana* accessions. They form connected graphs whose topography differs substantially among phenotypes. We focus our analysis on the incidence of epistasis (non-additive interactions among mutations) because a high incidence of epistasis can reduce the accessibility of evolutionary paths towards high or low phenotypic values. We find epistatic interactions in 67 phenotypes, and in 51 phenotypes every pairwise mutant interaction is epistatic. Moreover, we find phenotype-specific differences in the fraction of accessible mutational paths to maximum phenotypic values. However, even though epistasis affects the accessibility of maximum phenotypic values, the relationships between genotypic and phenotypic change of our analyzed phenotypes are sufficiently smooth that some evolutionary paths remain accessible for most phenotypes, even where epistasis is pervasive. The genotype network representation we use can complement existing approaches to understand the genetic architecture of polygenic traits in many different organisms.

**Author summary:** How genotypic change relates to phenotypic change is a fundamental problem in evolutionary biology. A smooth relationship between genotypic and phenotypic change facilitates phenotypic evolution, whereas a rugged one hampers it, because evolutionary change on a rugged landscape does not allow a monotonic increase or decrease in phenotypic value. We use data from multiple genome-wide association studies for 80 different phenotypes in the thale cress *Arabidopsis thaliana* to construct 80 genotype-phenotype maps for these phenotypes. We use a network representation for each map, where a vertex corresponds to a genotype, and where two vertices are connected if they differ in a single nucleotide. We study the incidence of non-additive interactions between mutations as a proxy for the ease to reach extreme phenotypic values of these maps. This incidence varies greatly among phenotypes, but even where it is highest, some monotonic pathways to high or low phenotypic values exist for the vast majority of phenotypes, such that the underlying phenotypes are accessible to adaptive evolution.

## Introduction

Understanding how genotypes are linked to phenotypes and ultimately to fitness is of central interest for evolutionary biology, because genotypic changes in the form of DNA mutations cause heritable variation in phenotypes, on which natural selection acts. A common approach to study the relationship between genotypes and phenotypes relies on the concept of genotype-phenotype maps introduced by Alberch [1]. Such maps have been characterized for phenotypes on multiple levels of biological organization. For example, these maps relate RNA genotypes to RNA phenotypes [2, 3], short amino acid sequences to folded proteins [4, 5], genotypes of regulatory networks to gene expression phenotypes [6, 7], genotypes of metabolic networks to nutrient consumption phenotypes [8, 9], and genotypes of developing organisms to the morphologies of macroscopic body structures like teeth [10]. Genotype-phenotype maps have even been characterized for digital organisms [11]. While many early studies used computational approaches, genotype-phenotype maps are increasingly studied experimentally, for instance in a transfer RNA [12], a ribozyme [13], a small nucleolar RNA [14], a green fluorescent protein [15], and the antibiotic resistance beta-lactamase TEM-1 [16]. Both empirical and experimental studies of such maps address fundamental evolutionary questions, for example about the implications of pleiotropy for the evolvability of complex organisms [17], the robustness and persistence of bet-hedging [18], the role of epistasis in evolution [19], or the influence of genetic correlations on mutational robustness and evolvability [20].

A genotype-phenotype map can be represented as a network of genotypes. The vertices (or nodes) of this network are genotypes, that is, nucleotide or amino acid sequences. Two genotypes are connected by an edge (that is, they are neighbors), if the corresponding sequences differ in a single small mutation, such as a nucleotide or amino acid change [2,21,22]. Genotype networks can provide insights into the role of evolutionary constraints for adaptive evolution [22], the role of convergence [23], or the evolution of binding affinities of transcription factors to DNA [24–26]. Moreover, they can help us understand a fundamental characteristic of biological entities, namely their evolvability. Evolvability describes the potential of a biological system to bring forth adaptive heritable variation, such as mutations that increase fitness [27, 28]. In genotype networks, a succession of such mutations constitutes an accessible path, that is, a path of edges along which fitness increases monotonically. Any one high fitness genotype may be reachable through multiple paths, but only some of these paths may be accessible, while others traverse fitness valleys. This fraction of accessible paths can serve as a proxy of evolvability, because it quantifies how easily high fitness genotypes can be reached through a series of mutations that never decrease fitness [25,29–33]. One important factor that can affect path accessibility is epistasis, that is, non-additive interactions between mutations, because it can create steps of decreasing fitness along a mutational path, and thus reduce path accessibility [28,34–39].

Here we apply the genotype network concept to experimentally determined organism-level phenotypes in order to study phenotypic accessibility and epistasis. Specifically, we characterized the architecture of genotype networks for 80 different quantitative phenotypes of the thale cress *Arabidopsis thaliana*, whose association with genotypes has been characterized in multiple genome-wide association studies [40]. A genome-wide association study (GWAS) identifies genomic sites at which nucleotide changes affect a phenotype. For any one genomic position, it quantifies whether different nucleotide variants are statistically associated with the phenotypic value of different individuals in a population. The method was pioneered in human genetics, where it is used to understand complex genetic traits that underlie different diseases [41]. In plants, a GWAS is often used to study the genetic architecture of agronomically important traits like crop yield and quality [42–46]. *Arabidopsis thaliana* is ideally suited for a GWAS because its natural inbred lines (called accessions) can be maintained via self-fertilization, such that many phenotypes of genetically similar individuals can be scored. Thus, multiple traits in *A*. *thaliana* have been successfully analyzed in a GWAS [40,47–54].

Our work builds on a pioneering study that performed a GWAS on 107 different phenotypes of *A. thaliana* [40]. Among these 107 phenotypes are 80 quantitative traits. We built a genotype network for each of these 80 phenotypes and characterized its structure. We report pervasive epistasis in these genotype networks. Even though such epistasis can reduce the accessibility of mutational paths to extreme phenotypic values, a substantial fraction of all mutational paths to each of our analyzed genotypes remains accessible.

## Results

### Building genotype networks whose size depends on the chosen P value cutoff

We first reconstructed separate genotype networks for each of the 80 quantitative phenotypes. These genotypes fall into four categories, namely flowering, development, defense, and the concentrations of various ions. Phenotypes in each category are highly diverse. For example, flowering phenotypes include the expression level of the MADS-box transcription factor *FLOWERING LOCUS C* that inhibits flowering until exposure to a prolonged cold period (that is, after vernalization [55]). They also include the number of days required for the formation of first buds at a growth temperature of 16°C. Developmental phenotypes include the germination rates of seeds that were stored for 28 days under dry conditions, but also the hypocotyl length after seven days of growth. Defense phenotypes include the number of colony forming units of the plant pathogen *Pseudomonas syringae* pv. *tomato* DC3000, and the number of offspring of the aphid *Myzus persicae*. Finally, the ion phenotype category comprises concentrations of two type of ions, those of heavy metals such as nickel or cobalt, and those of trace elements required for processes as diverse as stress resistance (*e.g.* potassium [56]) and enzyme catalysis (*e.g.* calcium [57]). We will discuss results for these four phenotypic categories separately throughout the paper.

The number of genomic positions associated with a phenotype depends on the *P*-value threshold below which a statistical association between phenotypic value and genotype at that position is deemed significant [58]. We first chose a specific *P*-value cutoff, and considered all genomic positions as associated with a phenotype below this cutoff as sites that affect the phenotype (Fig 1A). We then concatenated, separately for each accession, the nucleotides at each significant locus into a single nucleotide string. The length of this string is equal to the number of significant positions (Fig 1B). This string is a compact representation of the accession’s genotype or, more specifically, of its haplotype, at those genomic positions that affect the phenotype. Each unique string corresponds to one vertex in the genotype network. Two vertices are connected if they are nearest neighbors, that is, if they differ in exactly one nucleotide site (Fig 1C).

**Fig 1.**
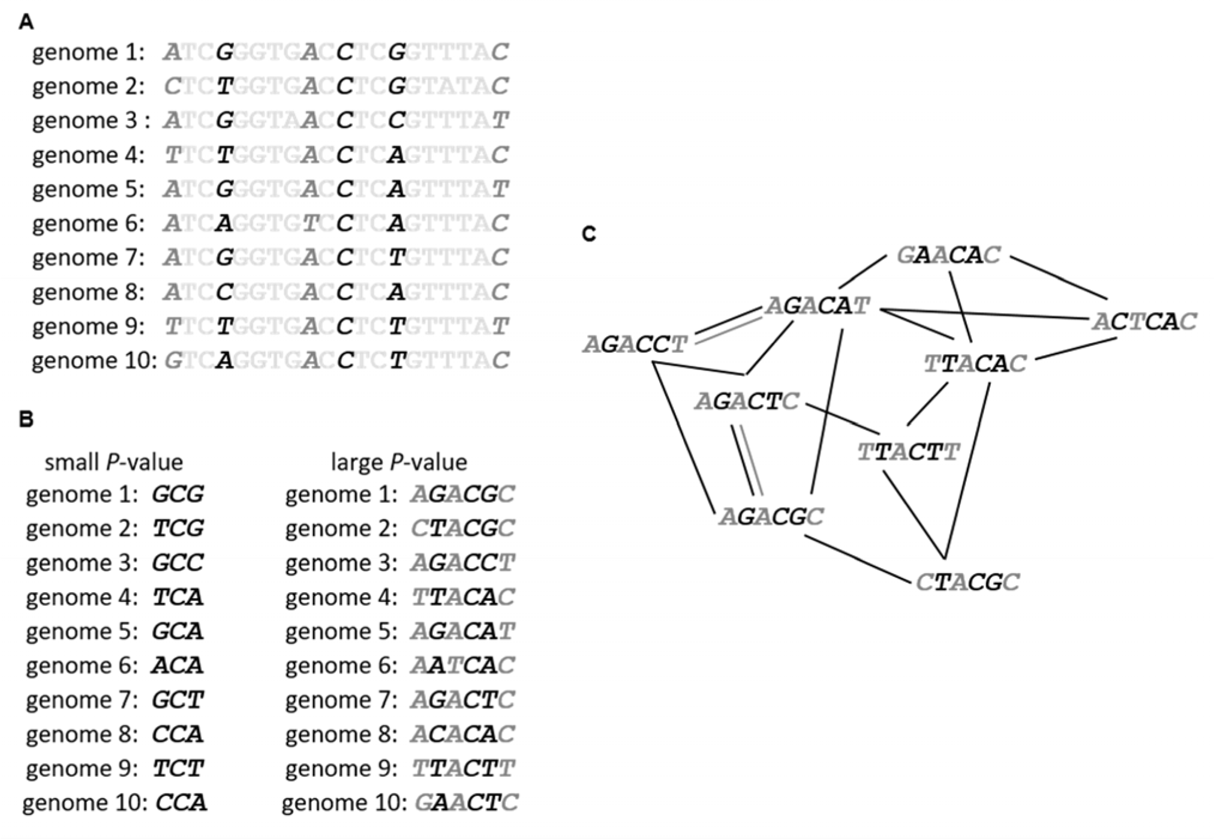
How to build a genotype network, illustrated with a hypothetical example. (A) Genome-wide association studies identify genomic positions affecting a phenotype. This statistical association is expressed with *P*-values, which are small (black type face in this hypothetical example) for genomic positions strongly associated with a phenotype. They are larger (dark grey type face) for genomic positions more weakly associated with the genotype. Light grey type face indicates nucleotides with no significant association to the phenotype. (B) Nucleotides at positions significantly associated with a phenotype are concatenated into one nucleotide string. The resulting string will be shorter if only strongly associated positions (smaller *P*-values) are considered (black letters, left hand side) and longer if also more weakly associated positions are considered (larger *P*-values, right hand side, black and dark grey letters). (C) Each nucleotide string of significantly associated positions becomes one vertex in a graph that we refer to as a genotype (haplotype) network. Two such vertices are connected by an edge if they differ in exactly one nucleotide. Black edges connect shorter nucleotide strings resulting from smaller *P*-values, and dark grey edges connect longer nucleotide strings resulting from larger (but still significant) *P*-values. We note that fewer pairs of nucleotide strings can be connected through edges as the length of nucleotide strings (and the *P*-value) increases. We note further that in biological data, multiple individuals may have the same genotype at all positions associated with the phenotype. In this case (not shown), we represent multiple individuals by only one network vertex. For didactic reasons, our hypothetical example uses multiallelic genome positions, whereas the data we analyze comprises only biallelic positions.

If we choose a large (non-stringent) *P*-value in this construction, many loci will appear to be associated with the phenotype, and the resulting nucleotide string will be long (Fig 1B). However, the likelihood that two such strings differ in exactly one position decreases with the length of the string. In consequence, most genotypes may be very different from one another, and only few genotypes will be neighbors. Such a genotype network will typically have many vertices, but it may be fragmented into multiple small connected components – sets of vertices that can be reached from each other through a sequence of edges. As this *P*-value decreases to become more and more stringent, each genotype will include fewer and fewer genomic positions, resulting in shorter nucleotide strings (Fig 1B). As these strings become shorter, it also becomes more likely that they become identical to each other, and the number of strings with one nucleotide difference will become smaller. Both cases – genotype strings that are too long (large *P*-values) or too short (small *P*-values) – are not conducive for network analysis. For each phenotype, we thus explored a broad range of *P*-values between 10^-4^ (largest published value and least stringent) and 10^-9^ (most stringent), to identify one that yielded a network with the maximum number of squares (that is, cycles of length four) and a maximum component size. We aimed to maximize the number of squares in a genotype network, because their presence is required for the analysis of epistasis (see Methods).

### Connected genotype networks can be built for whole-organism phenotypes

For all 80 phenotypes, we found that intermediate *P*-value cutoffs between 10^-4.5^ and 10^-6^ yielded genotype networks with the highest number of squares and the largest component size (S1 Fig, S1 Table, and S2 Table). In each of the four phenotype categories (flowering, ions, defense, development), a single *P*-value lead to one network with a maximum number of squares. The network component with the largest number of squares comprised 39 squares and 45 vertices (flowering phenotypes), 10 squares and 13 vertices (ion phenotypes), 58 squares and 52 vertices (defense phenotypes), and 27 squares and 36 vertices (developmental phenotypes; S1 Fig, S1 Table and S2 Table). Fig 2 shows the structure of these four genotype networks, which represent the following phenotypes: (i) in the flowering category, plant diameter at flowering (phenotype 1, abbreviated as ‘FT Diameter Field’ in [40], i.e., the largest diameter of the plant on the day where the first flower was observed, Fig 2A); (ii) in the ions category, the concentration of arsenic (phenotype 67, ‘As75’ in [40], Fig 2D); (iii) in the defense category, the number of colony forming units of the bacterial plant pathogen *Pseudomonas viridiflava*, to which we refer below as ‘bacterial growth’ (phenotype 70, ‘As CFU2’ in [40], Fig 2G); and (iv) in the development category, the diameter of plants grown at 16 °C at eight weeks after germination, which we refer to as ‘plant width’ (phenotype 14, ‘Width 16’ in [40], Fig 2J). *P*-values leading to the maximum number of squares in each genotype network did not necessarily yield the maximum component size in each network (S2 Fig and S2 Table). However, since we aimed to study the influence of epistasis on the evolvability of whole-organism phenotypes, we chose *P*-value cutoffs leading to the largest number of squares in a network component for subsequent analyses. The number of squares in the four example networks is comparable to random networks for all but the defense genotype network, where the number of squares is much higher in the biological data (S1 Text and S3 Fig). For most analyses below, we will present detailed results for the four genotype networks shown in Fig 2, and discuss summary statistics for the remaining 76 genotype networks. At our chosen *P*-value threshold 13 genome positions (loci) are associated with the flowering phenotype, 10 positions with the ions phenotype, 79 positions with the defense phenotype, and 24 positions with the development phenotype (S1 Table; see S3 Table for annotations of protein coding genes in which these genomic positions lie).

**Fig 2.**
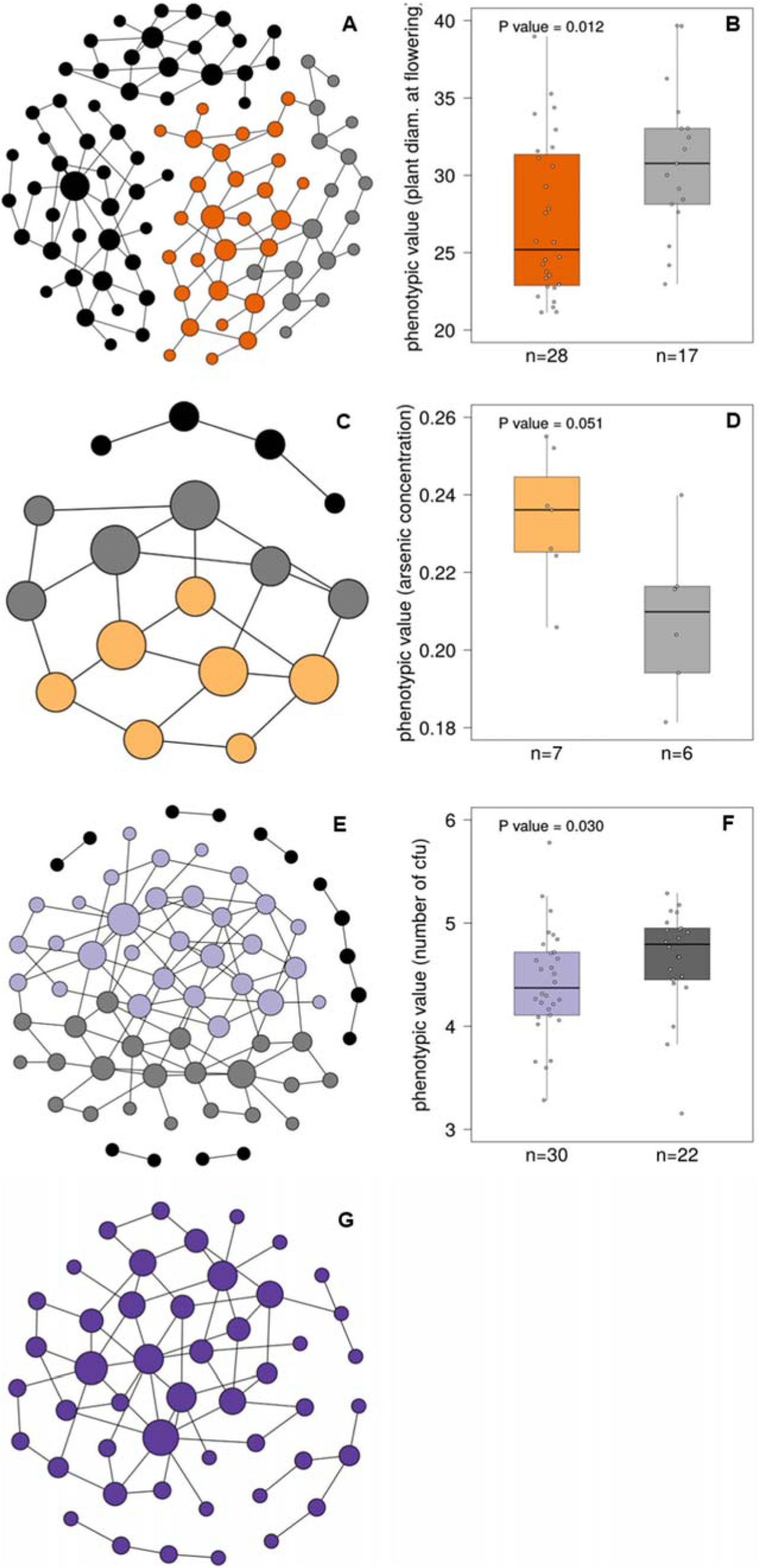
Genotype networks for four example phenotypes from each of the four phenotype categories. The four depicted networks correspond to the phenotypes plant diameter at flowering (panel A), arsenic concentration (panel C), bacterial growth (panel E), and plant width (panel G). In each network, a vertex (circle) represents a genotype (nucleotide string) and two vertices are connected by an edge if the underlying genotypes differ in one nucleotide. The size of each vertex is proportional to its vertex betweenness, which is the fraction of shortest paths through all vertex pairs that visit this vertex. Each network was laid out by the force-directed Fruchterman-Reingold graph embedding algorithm implemented in Gephi [59]. Networks shown in (A), (C) and (E) can be subdivided into two modules and each vertex in these networks is colored according to its module membership. Vertices of components that are not connected to the largest network component are colored black. Isolated vertices (that is, genotypes without a one-mutant neighbor) are not depicted. No modular structure could be detected in the genotype network of plant width (G). Box plots of panels (B), (D) and (F) show the phenotypic values in each module, where the box color is the same as that of the vertices in panels (A), (C), and (E), respectively. In each box plot, the central horizontal line indicates the median value, the lower and upper box limits correspond to first and third quartiles, whiskers indicate values within the 1.5-fold interquartile range, and each open circle shows one data point (vertex phenotype), illustrating the distribution of the phenotypic values. The number of vertices in each module is indicated as the sample size (n) on the horizontal axis. Phenotypic values differ significantly (Wilcoxon Rank-Sum Test) between genotypes in different modules of the two phenotypes plant diameter at flowering (panel B) and bacterial growth (panel F).

### Three of four largest genotype networks show a modular structure

Modularity is a common feature of biological networks [60–62], and it has also been reported for genotype networks, for example those of RNA molecules and the hemagglutinin sequences of influenza A viruses [3,23,63]. To identify modules (or ‘communities’) in our genotype networks, we applied an algorithm developed by Blondel *et al*. [64] as implemented in Gephi [59]. This revealed that three out of the four networks could be partitioned into two modules. The two main modules found in each network are highlighted by different coloring of vertices in a genotype network (Fig 2A, Fig 2C, and Fig 2E). Phenotypic values differed significantly between the two modules in the flowering and defense genotype network (Fig 2B and Fig 2F; *P-*values ≤ 0.03007, Wilcoxon Rank-Sum test). However, the biological significance of modularity is difficult to assess in our analysis, because our genotype networks are generally small. We also investigated the relation between the number of mutational steps that separate vertices in a genotype network and the phylogenetic distance between accessions, but found only a weak association between these quantities (S2 Text and S4 Fig). This is not surprising, because loci that affect any one phenotype may diverge at rates different from that of an entire genome, especially if the phenotype is subject to selection.

### Pervasive epistasis in genotype networks of whole-organism phenotypes

Because our analyses are based on genomic positions associated with phenotypic change [65] one would expect that neighbors in a genotype network – which differ at exactly one phenotype-associated position – have different phenotypic values. This may not be the case, however, if the phenotypic effects of mutations strongly depend on the genetic background in which they occur, that is, if epistasis is pervasive. To investigate the effect of each single mutational step in a genotype network, we first quantified the amount of phenotypic variation (δ) in our data that is due to non-genetic causes, such as measurement noise. Specifically, we computed for each phenotype the average coefficient of variation for phenotypic values measured from accessions where all phenotype-associated positions had the same genotype (Methods and S1 Table). We considered two phenotypic values as identical if they differed by less than δ.

Based on this criterion, many neighboring genotypes in a genotype network showed identical phenotypic values (Fig 3A). Motivated by these observations, we next quantified epistasis in a more fine-grained way by following a previously established approach [25]. This approach is based on analyzing individual squares in a genotype network (Fig 3B). It allows us to discriminate between magnitude epistasis, simple sign epistasis, and reciprocal sign epistasis (Fig 3C). In magnitude epistasis, the phenotypic value of a ‘double’ mutant is higher than the sum of the phenotypic values of two ‘single’ mutants (Fig 3C, subpanel *ii*). In simple sign epistasis, one ‘single’ mutant has a higher phenotypic value and the second ‘single’ mutant shows a lower phenotypic value compared to the ‘wild type’ (Fig 3C, subpanel *iii*). In reciprocal sign epistasis, both ‘single’ mutants have a lower phenotypic value than the ‘wild type’ (Fig 3C, subpanel *iv*). We quantified each of these three kinds of epistasis in each phenotype and grouped the results by our four major phenotype categories (Fig 3D).

**Fig 3.**
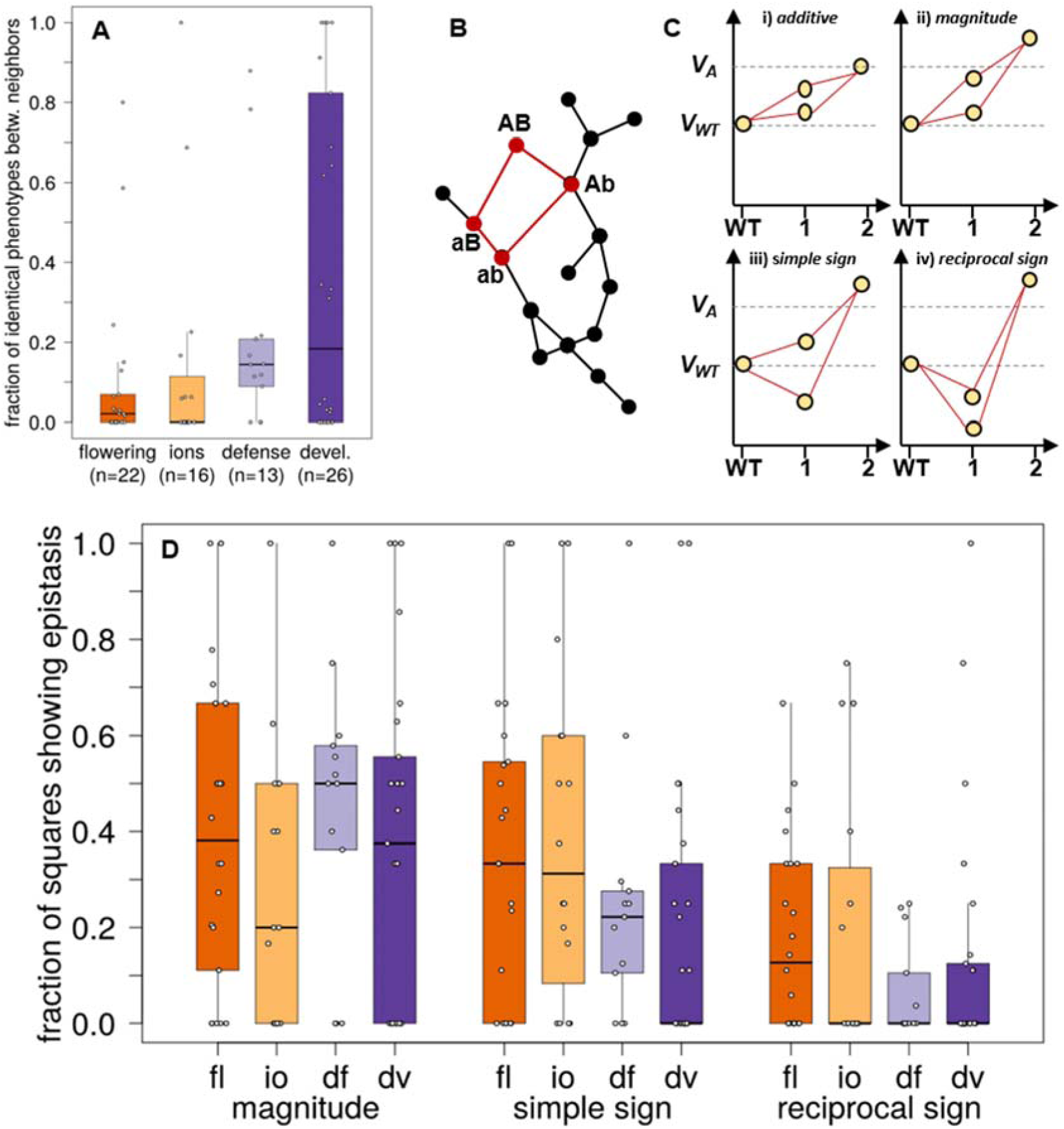
Pervasive epistasis in whole-organism phenotypes. (A) Fraction of genotype network neighbors with identical phenotypic values. We consider the absolute difference between the phenotypic values of two neighbors as identical if it does not exceed the coefficient of phenotypic variation for accessions with the same genotype at all phenotype-associated loci (Methods). We plot the fraction of neighbors with identical phenotypic values (vertical axis) in the form of a box plot for each of the four phenotype categories (“devl.”: development). The number of phenotypes in each category is shown on the horizontal axis as sample size (n). (B) Detection of epistasis. The presence of squares in a genotype network (highlighted in red) permits the analysis of epistatic interactions. Epistasis is detected by comparing the phenotypic values associated with the four vertices in a square, as shown in panel C. (C) We distinguish different types of epistasis. Vertical axes show phenotypic values in arbitrary units. V*_WT_* and V*_A_* denote the phenotypic value of the wild type and the expected phenotypic value in an additive (non-epistatic) scenario. The circles in each plot indicate a ‘wild type’ sequence, two of its one-mutant neighbors and the double mutant neighbor as indicated by the horizontal axes (WT, wild type). Panel i: the phenotypic values of the single mutant neighbors are higher than the wild type value, and their sum equals the phenotypic value of the double mutant (additive scenario). Panel ii: the phenotypic values of the single mutant neighbors is again higher than the wild type value, but the value of the double mutant is greater than the sum of the values of the single mutant neighbors (magnitude epistasis). Panel iii: the phenotypic value of one single-mutant neighbor is lower than the value associated with the wild type (simple sign epistasis). Panel iv: The phenotypic values of both single-mutant neighbors are lower than the wild type value (reciprocal sign epistasis). (D) Frequency of different types of epistasis. The vertical axis shows the fraction of squares with magnitude, simple sign, and reciprocal sign epistasis as a box plot, where data are grouped according to the type of epistasis and phenotype category (horizontal axis; “fl”: flowering; “io”: ions; “df”: defense; “dv”: development). In each box plot, the central horizontal line indicates the median value, lower and upper box limits show the values of the first and third quartile, whiskers indicate values within the 1.5-fold interquartile range, and each open circle shows one data point.

We found epistasis in 67 out of 76 investigated phenotypes, and in 51 phenotypes, each pairwise interaction was epistatic (S1 Table). Epistasis of all three kinds was abundant in each phenotype category, affecting up to 100 % of all pairwise interactions in a genotype network (S1 Table). For example, the genotype network of the phenotype days until flowering at 10°C (phenotype 31) contained 17 squares, of which 12 showed magnitude epistasis, four showed simple sign epistasis, and one showed reciprocal sign epistasis. In the ion category, networks of the phenotypes magnesium concentration and selenium concentration (phenotypes 105 and 33) contain eight squares, and each of them showed magnitude or simple sign epistasis. The defense phenotype bacterial disease resistance (phenotype 42) was also found to be highly epistatic. In nine squares, we found five with magnitude epistasis, two with simple sign epistasis and two with reciprocal sign epistasis. Finally, the phenotype days to germinate at 10°C (development; phenotype 59) contained 14 squares among which 12 showed magnitude epistasis and two showed reciprocal sign epistasis. Moreover, magnitude epistasis occurred in all six squares of the developmental phenotype hypocotyl length (phenotype 44), and we detected simple sign epistasis in all four squares of the developmental phenotype number of days between germination and senescence of the last flower (phenotype 58). In contrast, we found no epistasis in the two defense phenotypes bacterial disease resistance (phenotypes 103 and 83), the two ions phenotype copper and arsenic concentration (phenotypes 24 and 67), and in five developmental phenotypes, namely vegetative growth during and after vernalization (phenotype 72 and 39), secondary seed dormancy after six weeks of cold treatment (phenotype 106), seedling growth rate (phenotype 90) and lesioning measured by dry weight (phenotype 93) (S1 Table).

We also detected a high frequency of epistatic interactions when we chose a different *P*-value cutoff for our network reconstructions (S5 Fig). Specifically, we used for this sensitivity analysis *P*-values that are 0.5 scores (-log_10_) higher or lower than *P*-values leading to genotype networks with the largest number of squares. Moreover, we detected also a high fraction of epistatic interactions when we took potential linkage disequilibrium into account (S3 Text and S6 Fig), and when we ignored genomic positions with rare minor alleles (frequency below five percent; S7 Fig).

### Accessibility of evolutionary paths to minimum and maximum phenotypic values

A genotype network represents a sample of genotypes from an adaptive landscape, and to study such landscapes, it is useful to determine the number of fitness peaks and the number of accessible (fitness-increasing) paths to these peaks. Our phenotype data do not contain direct measurements of fitness, with the possible exception of fitness-proxies, such as a low number of colony forming units of plant pathogenic bacteria (phenotypes 21 and 70) as indicators of high pathogen resistance. However, even the number of mutational paths to maximal (minimal) phenotypic values along which a phenotype monotonically increases (decreases) can be informative about evolutionary dynamics [25,66,67], because a higher fraction of accessible mutational paths may reflect higher evolvability [28].

To find out how accessible minimum and maximum phenotypic values are for all of our genotype networks, we first enumerated the distribution of the length of all paths through the genotype network associated with each phenotype. Unsurprisingly, this distribution strongly depends on the phenotype (S8 Fig). We then focused only on those paths that reach the phenotype with the highest phenotypic value and determined the fraction of paths that monotonically increase the phenotypic value. We refer to these paths as mutationally accessible [29–31]. To ensure comparability of results across phenotypes, we considered only genotype networks where a single vertex is associated with the maximum phenotypic value. Below, we only discuss the fraction of accessible paths to the maximum value in detail, because analyzing the fraction of accessible paths to the phenotypic minimum yielded similar results (S9 Fig). We present results separately for each phenotype category (Fig 4).

**Fig 4.**
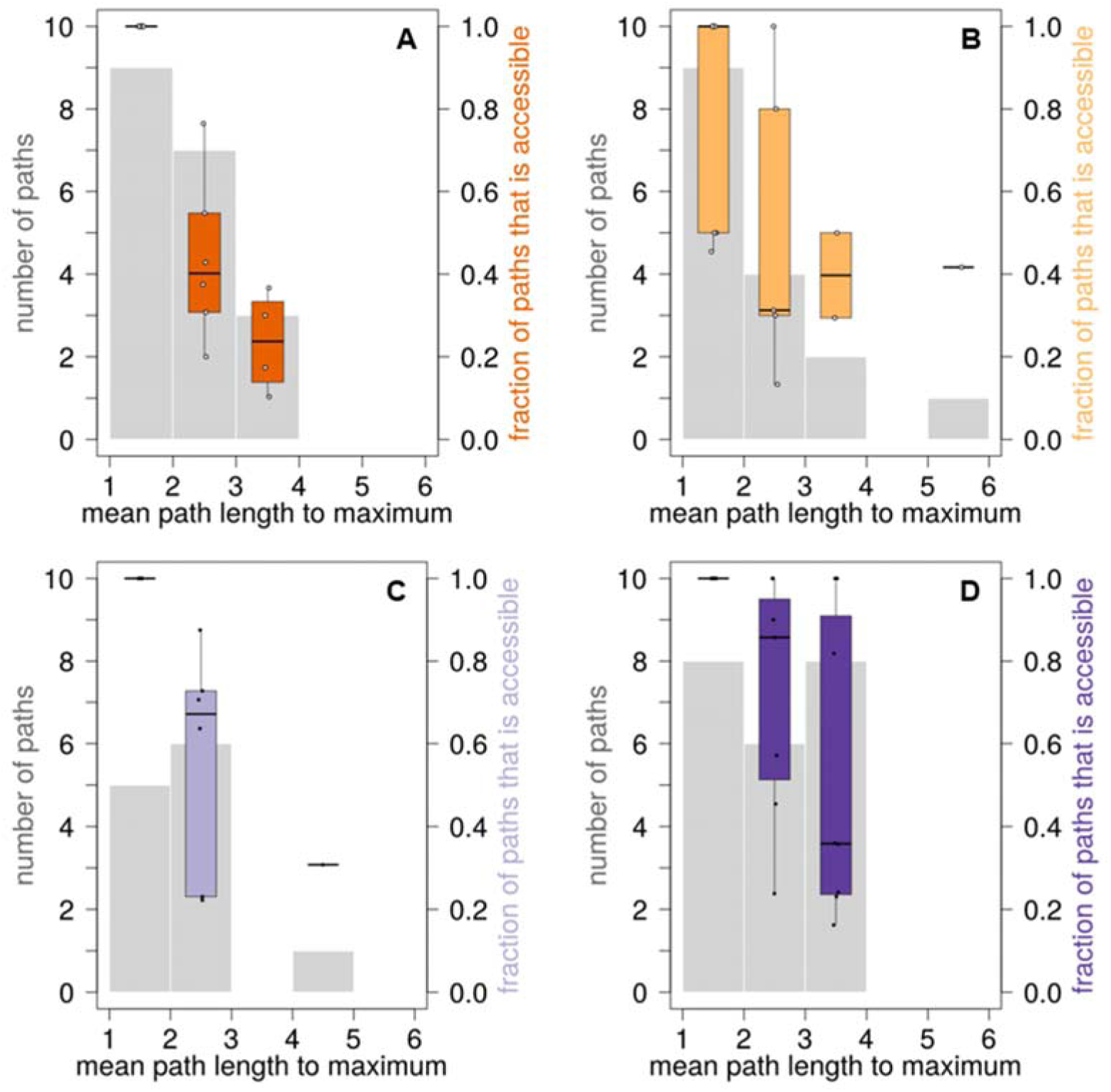
Distribution of mutational path lengths to maximum phenotypic values and fraction of accessible paths. For each phenotype, we calculated the mean length of the mutational path from each vertex in the network to the vertex with the maximum phenotypic value. We show the number of paths (left vertical axes) binned according to path length (horizontal axes, path length between 1 and 2 mutational steps, between 2 and 3 mutational steps, etc.) as indicated by the grey histograms. The colored box plot in each bin shows the fraction of paths to the maximum phenotypic value that is accessible (right vertical axes). Each panel shows the result for one phenotype category, namely flowering (A), ions (B), defense (C), and development (D).

The fraction of accessible paths generally decreased with increased path length, but the rate of this decrease differed between phenotypes. For example, all mutational path with a length between 1 and 2 were accessible in all flowering phenotypes (Fig 4A), in all defense phenotypes (Fig 4C), and in all developmental phenotypes (Fig 4D), but not in all ion phenotypes (Fig 4B). Specifically, in the ion phenotype boron concentration (phenotype 54), only 45.5 % of such mutational paths were accessible (S1 Table), and in the ion phenotypes potassium and sulfur concentration (phenotypes 81 and 91) 50 % of paths were accessible. The maximum fraction of accessible mutational paths with a length between two and three assumed its lowest values of 13.3 % in the ion phenotype calcium concentration (phenotype 51), and 20.0 % in the flowering phenotype days to flowering at 22°C (phenotype 102). It assumed its highest values of 100 % in both the ion phenotype nickel concentration (phenotype 64), and in the developmental phenotypes days to germination at 22°C (phenotype 45) and vegetative growth rate (phenotype 72). Together, these data illustrate that the ability to increase a phenotype monotonically strongly depends on the phenotype.

Again, we performed a sensitivity analysis for these results. While the length of mutational paths to maximum phenotypic values decreases in smaller networks, a large fraction of mutational paths remains accessible if we construct networks with higher or lower *P*-value cutoffs (S10 Fig). We conducted this analysis for *P*-values that vary by one order of magnitude around the default P-value yielding genotype networks with the largest number of squares (S10 Fig). We also found a high accessibility of mutational paths to maximum phenotypic values when considering the effect of linkage disequilibrium (S3 Text and S6 Fig). The same holds when we ignored genomic positions with rare minor alleles (frequency below five percent) when building genotype networks (S7 Fig).

One factor that influences the fraction of accessible mutational paths is the incidence of sign epistasis, which can render a mutational path inaccessible (Fig 3C and [28, 68]). The presence of epistasis strongly affects the fraction of accessible mutational paths in some of the phenotypes we studied. One class of noteworthy examples comprises phenotypes with short mutational paths and nonetheless low accessibility. A case in point is the ion phenotype Boron concentration (phenotype 54), where the average path to the maximum phenotypic value requires 1.9 mutations, but where only 45 % of paths are accessible (S1 Table). Consistent with an expectation of substantial epistasis, three out of four squares in the genotype network for this phenotype showed reciprocal sign epistasis and one square showed simple sign epistasis (S1 Table). Another example is the ion phenotype calcium concentration (phenotype 51), where paths to the maximum phenotypic value have a mean number of 2.67 mutational steps, but where only 13.3 % of these paths are accessible (S1 Table). For this phenotype, we found reciprocal sign epistasis in four out of six squares, as well as simple sign epistasis and magnitude epistasis in the remaining two squares (S1 Table). The prevalence of reciprocal sign and simple sign epistasis limits the number of accessible paths.

Conversely, a low incidence of epistasis can render even long paths to some phenotypes highly accessible. Among the most extreme examples is the developmental phenotype primary dormancy after seven days of dry storage (phenotype 20). Here, an average of 3.06 mutational steps are needed to reach the maximum phenotypic value, but 100 % of all paths are accessible. In the phenotype vegetative growth rate after vernalization (phenotype 39), an average of 3.4 mutational steps are required to arrive at the maximum phenotypic value, and again, 100 % of all paths are accessible. The reason is that epistasis is largely absent in both phenotypes, with the exception of one square that shows magnitude epistasis in the latter phenotype (S1 Table). Of note is also the defense phenotype bacterial disease resistance (phenotype 2). Even though it has a long average path of 4.62 mutational steps to the maximum phenotypic value, almost a third (30.8%) of these paths are accessible (S1 Table). The reason is again a rarity of epistasis: 14 out of 27 squares in its genotype network show no epistasis, and four squares show only magnitude epistasis, which does not affect path accessibility (S1 Table). Similarly, in the ion phenotype magnesium concentration (phenotype 105), the average mutational path length is 5.25 steps, 41.7 % of all paths are accessible (S1 Table), and none of its squares shows reciprocal sign epistasis.

When considering all phenotypes, we detected a negative correlation between the incidence of reciprocal sign epistasis and the fraction of accessible mutational paths (Kendall’s tau = −0.1973; *P*-value = 0.04141; n = 69). This result is consistent with previous studies demonstrating that reciprocal sign epistasis limits the accessibility of mutational paths [28, 68]. The association between simple sign epistasis and the mean fraction of accessible mutational paths was also negative but not significant (Kendall’s tau = −0.154; *P*-value = 0.0943; n = 69), which we attribute to our modest sample size, and to the fact that simple sign epistasis reduces path accessibility less than reciprocal sign epistasis (Fig. 3C). Also, we observed no significant correlation between magnitude epistasis and the mean fraction of accessible mutational paths (Kendall’s tau = −0.125; *P*-value = 0.1782; n = 69), which is consistent with the fact that magnitude epistasis does not affect path accessibility (Fig. 3C).

## Discussion

We constructed genotype networks of whole-organism phenotypes from genome-wide association data for 80 quantitative *Arabidopsis thaliana* phenotypes that fall into four categories (flowering, ions, defense, and development). Representative networks from each category formed smaller or at most equally large connected components than networks constructed from randomized genotype strings with similar genetic diversity (S3 Fig). However, because our analysis is based on a limited number of 199 genotypes (accessions), the size of this component was generally modest and comprised only up to 115 genotypes. This modest size may help explain why network statistics like maximum vertex betweenness were not distinguishable from that of random networks in two of the four studied genotype networks.

The genotype networks we analyzed reveal that epistasis is widespread for organism-level phenotypes. Such epistasis has also been observed for diverse types of molecular phenotypes, for example those of the green fluorescent protein [15], the beta-lactamase TEM-1 conferring resistance to ampicillin [69], a tRNA gene [12], or in the binding of transcription factors to DNA [25]. Epistasis has also been reported for organism-level phenotypes of *A. thaliana*, for example for metabolomics phenotypes [70], stem branching [71], or flowering in different environments [72]. However, these studies rely on quantitative trait loci (QTL)-mapping, which provides associations between genomic regions and phenotypes. In contrast, our analysis of epistasis is based on mutations in single genomic positions and thus allows a fine-grained analysis of epistatic interactions that involve single nucleotide changes. In addition, our analysis is built on many (80) and very different kinds of quantitative phenotypes. It shows that epistasis is widespread for organism-level phenotypes and pervasive for some phenotypes, affecting *all* pairwise genetic interactions in 51 phenotypes (S1 Table; we note that in nine of these cases, only one square was present in the genotype network). Together with previous work, our analysis underscores that epistasis affects all levels of biological organization.

Pervasive sign epistasis influences phenotypic evolvability, because it can render mutational paths to extreme phenotypes inaccessible [28, 68]. When quantifying the accessibility of mutational paths for each phenotype, we found that it strongly depends on the phenotype. For example, even short paths of one to two mutational steps were inaccessible in the three ion phenotypes potassium concentration, sulfur concentration, and boron concentration. Conversely, in the ion phenotype magnesium concentration, 41.7 % of all paths between five and six mutational steps were accessible. While we found epistasis in almost all studied phenotypes (67 out of 76), it did not prevent the accessibility of maximum phenotypic values for the vast majority of phenotypes. That is, in 69 analyzed phenotypes, at least 10.3 percent of paths to the maximum were accessible.

We note that at least two factors beyond the scope of our work may help increase the accessibility of phenotypes we study here. First, because of the small number of accessions and the resulting small genotype network sizes, we did not consider epistatic interactions between more than two genomic sites, which can open paths that appear inaccessible when only pairwise interactions are considered [73–76]. Second, even strong and pervasive epistasis may not prevent phenotypic accessibility whenever populations are small and genetic drift is strong. The effective population size of *A. thaliana* is thought to be small, which may be explained by a strong bottleneck during the last ice age and a high frequency of inbreeding [77, 78]. Strong drift can help traverse fitness valleys created by epistasis, and thus render otherwise inaccessible paths accessible [79].

Our small sample size of less than 200 accessions causes further limitations of our study. Phenotypes influenced by rare genetic variants, or by genomic loci with small effects can only be detected in larger samples [80], such as a collection of 1,135 recently genotyped (but not yet as thoroughly phenotyped) *A. thaliana* accessions [81]. A GWAS with more accessions would also permit the identification of many more loci associated with a truly polygenic trait [58]. For our genotype network analysis, such larger sample sizes would have several consequences. First, because *A. thaliana* genomes are not highly polymorphic [81], multiple accessions in our genotype networks had identical genotypes at the loci associated with a given phenotype. A larger sample would increase the diversity at those positions, and it would lead to larger connected genotype networks. Larger genotype networks could also strengthen the tentative evidence for modularity we found in our modest data set, which would permit assessing the biological significance of this modularity. Moreover, larger genotype networks may have multiple vertices associated with high phenotypic values, which could further increase the accessibility of paths to extreme phenotypes. Finally, much greater sample sizes could help in analyses that were completely beyond the scope of this work. For example, paths through a large genotype network may affect multiple phenotypes, such that genotype network analysis may help understand not just pleiotropic effects of mutations, but how trade-offs and synergisms between multiple phenotypes affect a phenotype’s evolvability.

## Materials and Methods

### Data resources

We based our analysis on previously conducted genome-wide association studies in *A. thaliana* [40], where 199 accessions were phenotyped in 107 experimental conditions, and genotyped with a resequencing microarray covering 248,584 genomic positions. The genomic positions for this customized array were chosen on the basis of genomic polymorphism and linkage disequilibrium data [82]. For each of these phenotypes and genomic positions, the null hypothesis of no statistical association between phenotype and genotype was tested, and those genotypes significantly associated with a phenotype based on the rejection of the null hypothesis at a specific *P*-value were recorded. We restricted our analyses to all quantitative phenotypes from this study, of which there are 80 (S1 Table). These phenotypes can be grouped into four categories. They comprise phenotypes related to flowering (23 phenotypes), plant development (26 phenotypes), defense (13 phenotypes), and the concentrations of various ions in the plant (18 phenotypes). These ions include heavy metals, but also trace elements like potassium and calcium (S1 Table). The latter elements are, for example, necessary as enzyme co-factors or for stress responses. For each of the quantitative phenotypes, we retrieved the observed phenotypic values for each *A. thaliana* accession from the database AraPheno [83]. We also retrieved the genomic positions significantly associated with each phenotype, together with the corresponding *P*-value from the AraGWAS Catalog [84, 85]. This database contains recomputed GWAS results based on the imputed genotypes of the 1001 Genomes Consortium [81]. We obtained data for all analyzed genomic positions, that is, we did not exclude positions with rare alleles in our main analyses. However, to assess the sensitivity of our findings to the presence of genomic positions with minor allele counts, we removed genomic positions with minor allele counts lower than five percent (the default setting of the AraGWAS Catalog). Results for this analysis are summarized in S4 Table and S5 Table. We note that not all 199 accessions were used in each association study (S1 Table).

### Reconstruction and analysis of genotype networks for each phenotype

We first reconstructed separate genotype networks for each phenotype. To this end, we first chose a specific *P*-value cutoff, and considered all genomic loci associated with a phenotype below this cutoff as loci that significantly affect the phenotype. For the analyses described below, we varied these *P*-value cutoffs between 10^-4^ and 10^-9^, and note that a large *P-*value implies that more genomic positions are considered significantly associated with a phenotype (Fig 1B). We then concatenated, separately for each accession, the nucleotides at each significant locus into a single nucleotide string. The length of this string is equal to the number of significant positions (Fig 1B). This string is a compact representation of the accession’s genotype or, more specifically, of its haplotype, at those positions significantly associated with the phenotype. The number of these strings is equal to the number of accessions used in the association study for this phenotype. Our approach ensures that all resulting nucleotide strings are of the same length, and that the strings represent the same genomic positions, that is, they are aligned. For all resulting nucleotide strings that were identical among several plant accessions, we retained only one concatenated sequence, and used the mean phenotypic value across all accessions with this genotype string as the phenotypic value associated with this genotype. We call two nucleotide strings one-mutant neighbors if they differed in exactly one position. In all subsequent analyses, we considered only genotype strings that had at least one such neighbor. In the language of graph theory, this means that we analyzed only connected components of a genotype network that comprised at least two vertices [86].

We then used the nucleotide strings that had at least one neighbor as input for the R package igraph [87] to build a network in which each vertex corresponds to one of the strings, that is, to one genotype whose constituent nucleotides are significantly associated with the phenotype. Neighboring vertices correspond to neighboring strings, that is, to those genotype that differ in a single nucleotide. For each phenotype, we used multiple *P*-value cutoffs ranging between 10^-4^ and 10^-9^, that is, we constructed a genotype network for each of these cutoffs, and determined the number of connected components comprising at least two vertices, as well as the size of the largest component, to determine the *P*-value that yields a network best suited for network analysis. To identify modules in our network, we employed the algorithm of Blondel and colleagues [64] as implemented in Gephi version 0.9.2 [59]. We also used Gephi to display genotype networks.

### Creation of random genotype networks

To assess whether properties of the four example genotype networks presented in Fig 2 could be observed by chance alone, we performed randomization tests based on random genotype networks. Specifically, for each of the four example phenotypes plant diameter at flowering, zinc concentration, bacterial growth, and plant width, we created a random genotype network as follows. First, we determined the proportion of each of the four nucleotides in each column of the aligned genotype strings that we had used in creating the network. Second, we generated random nucleotide strings of the same length and with the same nucleotide composition provided by the GWAS data. Specifically, for each position of such a random string, we randomly chose a nucleotide with a probability equal to the proportion of this nucleotide in the corresponding column of the aligned genotype strings. We then built a genotype network for the resulting random strings in the same manner as for the biological genotypes. We repeated this procedure 10,000 times, and created one genotype network from each of the resulting 10,000 sets of random genotype strings. Similar to the analysis of networks based on biological data, we determined the number of squares in the resulting genotype network, the maximum component size, the maximum vertex betweenness, and the number of unique genotypes for each random genotype network.

### Reconstruction of phylogenetic relationships

To infer phylogenetic relationships between all 199 *A. thaliana* accessions, we employed the nucleotides at all 214,052 polymorphic positions present in the resequencing microarray [40]. Since each position on the array corresponds to one (polymorphic) genomic position, we concatenated the genotyped nucleotides for each accession into one contiguous nucleotide string. The strings from different accessions are of equal length (there were no missing data), which obviates the need for aligning them. We then used FastTree version 2.1.10 SSE3 [88] under a generalized time-reversible model (option-gtr) to construct a phylogenetic tree from these sequences with 1,000 bootstrap replicates. The resulting tree is illustrated in S11 Fig and the tree’s structure is provided in Newick format in S4 Text. We considered the number of internal nodes separating two accessions as a measure for the phylogenetic distance between them. We determined this number with the function distTips of the R package adephylo [89]. As a complementary distance measure, we also computed the number of nucleotide differences between all pairs of concatenated nucleotide sequences. To find out if one-mutant neighbors – genotypes differing in one nucleotide – in a genotype network also tend to be neighbors in the phylogenetic tree, we examined all pairs of neighboring genotypes in each genotype network and quantified how distant they are in the tree. Whenever one or both neighboring vertices in a genotype network were associated with multiple identical genotypes, we calculated the mean phylogenetic distance (number of intervening internal nodes and pairwise nucleotide differences) between *all* underlying genomes.

### Estimating measurement noise for quantitative phenotypes

To analyze how strongly phenotypes vary among different genotypes in a genotype network, it was necessary to estimate the non-genetic phenotypic variation for each phenotype. Since data from replicate experiments were only available for five phenotypes, we based this estimate on those accessions that had identical genotypes at all loci significantly associated with the phenotype. If the focal phenotype varies among such accessions, the cause is likely to be non-genetic. Specifically, for each of our 80 phenotypes, we identified all vertices in the associated genotype network for which at least three accessions had identical nucleotide sequences. For each such vertex, we then calculated the coefficient of variation of the phenotypic values for these accessions, that is, we divided the standard deviation of the phenotypic values by the mean of these values. Finally, we averaged the coefficients of variation for all vertices and used this average as measure of non-genetic phenotypic variation (δ) for the focal phenotype (S1 Table). Between 2.94 % and 80.0 % of all vertices in a network contributed to the calculation of δ (S1 Table).

### Detection of epistasis

To identify the presence of epistasis and to quantify the incidence of different types of epistasis, we first detected all squares in each genotype network by following a previously established approach [25, 90]. A square consists of four genotypes (vertices), which comprise a focal genotype, two of its one-mutant neighbors, and the double mutant that can be formed from the single mutants (Fig 3B and Fig 3C). By convention, we designated the vertex with the highest phenotypic value as the ‘double mutant’ genotype AB, and we ignored all squares where more than one vertex had this maximal value to ensure meaningful comparisons between squares. We called the ‘wild type’ genotype ‘ab’ and the remaining vertices ‘Ab’ or ‘aB’. Without epistasis (that is, when only additive interactions between genotypes occur), the phenotypic values *V* of the four genotypes should show the following relationship:

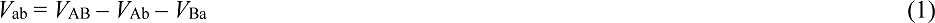

where *V*_AB_ is the phenotypic value of the double mutant, *V*_ab_ is the phenotypic value of the ‘wild type’ focal genotype and *V*_Ab_ and *V*_aB_ are the phenotypic values of the single mutants. If epistasis occurs, its magnitude can be calculated as follows:

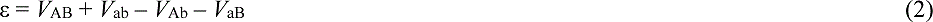

We considered epistasis to be present only if the absolute value of ε exceeded the threshold of non-genetic phenotypic variation (δ), whose calculation we described in the preceding subsection. We were able to calculate a value for δ for 79 of our 80 phenotypes (see above). For all squares of genotypes where ε < δ, we set the value of ε to zero, thereby allowing a conservative estimation of epistasis. Whenever epistasis was present (ε ≥ δ), we distinguished between magnitude, simple sign, and reciprocal sign epistasis ([30] and Fig 3C) as follows:

Magnitude epistasis:

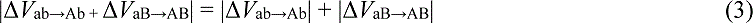

where Δ*V* denotes the difference in phenotypic values caused by one mutational step, for instance ab→Ab.

Sign epistasis: 

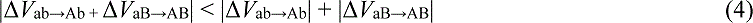

Reciprocal sign epistasis: 

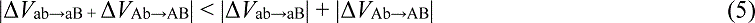

We performed this analysis only for 76 phenotypes, because the genotype networks for the two ion phenotypes molybdenum and cobalt concentration (phenotype numbers 73 and 22), as well as for the flowering phenotype number of leaves when the bolt reached five centimeters (phenotype number 36) contained no square. Moreover, the genotype network of the development phenotype days to germinate at 16°C (phenotype number 60) did not contain a square with a single maximum phenotypic value.

### Accessibility of minimum and maximum phenotypic values

We asked how accessible maximum (and minimum) phenotypic values are for all of our genotype networks. Specifically, we enumerated for each phenotype all paths through the associated genotype network that monotonically increase (decrease) the phenotypic value and that reach the genotype with the highest (lowest) phenotypic value. We refer to these paths as mutationally accessible [29–31]. We followed a previously established strategy [25] to compute this path. Briefly, we define a mutational path as the smallest number of edges (shortest path) that connects two vertices *i* and *j*. We designated the vertex with the highest (lowest) phenotypic value in a genotype as vertex *j*, iterated through all other vertices *i*, computed the shortest path between *i* and *j*, and identified for each path whether it is accessible, that is, whether the phenotypic value monotonically increases (decreases) along the path’s edges. Whenever we found several mutational paths with the same length between any two vertices *i* and *j*, we computed the accessibility for each path. We then calculated the fraction of accessible mutational paths, and refer to this fraction as the accessibility of minimum and maximum phenotypic values, respectively. In all these calculations, we considered the phenotypic values of two neighboring vertices as identical if the absolute difference of them does not exceed the coefficient of phenotypic variation (δ) for accessions with the same genotype at all phenotype-associated loci.

## S1 Text: Characteristics of genotype networks based on random nucleotide strings

To evaluate if properties of the four example genotype networks illustrated in Fig 2 would be expected to occur by chance alone for data sets of similar size and genetic diversity, we created 10,000 random genotype networks for each of the four phenotypes, computed network descriptors for them, and compared these descriptors with their values in the four biological networks. We built these 10,000 random networks from 10,000 sets of randomized genotype strings with the same nucleotide composition as the biologically observed genotypes (see Methods). The number of unique genotype sequences that this randomization procedure yielded was consistently higher than in the biological data for all 10,000 sets of strings and for all four phenotype categories (S3A Fig). In other words, the biologically observed genotype sequences are less diverse than expected by chance. One possible reason is that the corresponding genomic positions are under selection, because they are strongly associated with the investigated phenotype and thus functionally important. The number of squares in the genotype networks based on biological data lay within the range of the number of squares in the random genotype networks, with the exception of the defense phenotype bacterial growth, where the genotype network based on biological data showed a higher number of squares (S3B Fig). Moreover, the size of the largest network component of the flowering phenotype plant diameter at flowering and the defense phenotype bacterial growth lay within the range of the values obtained from random networks. In contrast, the largest component size was lower in the biological networks compared to random networks for the ion phenotype arsenic concentration and the developmental phenotype plant width (S3C Fig). We also determined the maximum vertex betweenness, that is, the highest number of shortest path that pass through a vertex (S3D Fig). It showed that the vertex betweenness of genotype networks based on biological data is in the lower range of those values obtained from random networks. For two phenotypes – the ion phenotype arsenic concentration and the development phenotype plant width – vertex betweenness was lower than in the random genotype networks, showing that vertices with very high betweenness can occur in random genotype networks of the size we study here.

## S2 Text: Mutational path length is weakly correlated to phylogenetic distance

The greater the distance is between two vertices in a genotype network, the greater is the number of mutations at phenotype-associated genomic positions that are needed to reach one vertex genotype from the other. We refer to this number of mutations as mutational path length. This length is not necessarily related to the phylogenetic distance between two accessions, because two accessions might be phylogenetically very different yet have very similar genotypes at the positions determining any one phenotype, indicating conservation of that genotype. Conversely, they may be phylogenetically very closely related, but have diverged greatly at the genomic positions affecting a focal phenotype, possibly indicating rapid evolution of the phenotype. To characterize the relationship between phylogenetic distance and mutational path length, we constructed a phylogenetic tree based on all 214,051 polymorphic loci of the 199 *A. thaliana* accessions used by Atwell and colleagues (2010). We used two measures of phylogenetic distances, namely the number of internal nodes (branch-points) that separate two accessions on the phylogenetic tree (S4A Fig), and the pairwise nucleotide distance between two accessions. For both measures, we found only weak correlations between the length of the shortest path between two accessions in a genotype network and phylogenetic distance in the four example phenotypes plant diameter at flowering, arsenic concentrations, bacterial growth, and plant width (S4B Fig and S4D Fig). More generally, mutational path length and the number of internal nodes (number of nucleotide differences) were weakly correlated for 26 (28) phenotypes (S4C Fig and S4E Fig, respectively). For the remaining phenotypes, we could not detect significant correlations.

Below, we discuss examples for two classes of noteworthy cases. First, two accessions may be neighbors in a genotype network (that is, they differ by one nucleotide in genomic positions that affect a phenotype), but they may nonetheless be phylogenetically distantly related, as quantified by the number of internal nodes or nucleotide differences between them. Second, two accessions may be closely related in phylogenetic terms, but separated by a long mutational path in a genotype network. Examples for the first scenario comprise the phenotypes sodium and molybdenum concentrations (phenotype 5 and 73) as well as leaf necrosis (phenotype 23). A single edge in the genotype networks of these three phenotypes connected one pair of accessions that are single-mutant neighbors, but that are phylogenetically separated by 27, 25.3 and 24.8 internal nodes, respectively (across all pairwise comparisons, the maximum number of internal nodes separating two accessions was 28). When using the number of nucleotide differences as a phylogenetic measure, we found that the genotype networks of the phenotypes days to germination at 10°C (phenotype 59), bacterial disease resistance (phenotype 70), and leaf number when the bolt reached 5 cm (phenotype 36) contained a single edge that connects two accessions as one-mutant neighbors, but the pair of accessions constituting this edge differed by 86,054 nucleotides, 85875.6 nucleotides, and 84,831.05 nucleotides, respectively. The maximum number of nucleotide differences between all pairs of accessions is 94,561.

Examples for the second scenario (close phylogenetic relations, but long mutational path between accessions) include the phenotype leaf number at flowering time (phenotype 41), plant width at 16°C (phenotype 14), and number of days between appearance of the first flower and senescence of the last flower (phenotype 32). The genotype network of these phenotypes contained accessions that are separated by five or more mutational steps in the genotype network, but only by one internal node in the phylogenetic tree.

## S3 Text: Effect of potential linkage disequilibrium on epistasis and path accessibility

One challenge in the analysis of GWAS data is the difficulty to discriminate between ‘causal’ genomic positions (loci) – those that affect a phenotype directly – and genomic positions that are only in high linkage disequilibrium (LD) with causal positions (Korte and Farlow, 2013). Atwell and colleagues (2010) reported that LD is usually too widespread to directly identify causal positions with a GWAS alone. We wanted to find out whether LD substantially affects the incidence of epistasis and the fraction of accessible mutational paths to maximum phenotypic values. Previous studies reported different degrees of LD in *A. thaliana*, and estimated LD to decay within 250 kbp (Nordborg *et al*., 2002), 10 kbp (Kim *et al*., 2007), and 5 kbp (Gan *et al*., 2011). The difference between these results originates from different methods to estimate LD, and from using global or local *A. thaliana* populations. To be conservative in our analysis, we assumed that any two genomic positions that are less than 250kbp apart might be in LD. We screened the genotype strings that we used to build genotype networks (at *P*-values that yield the largest number of squares) for loci within 250 kbp of each other. We found such positions in the genomic data for 60 phenotypes (S6 Table). Depending on the phenotype, we identified between zero and seven groups of genomic positions within 250 kbp of each other, and each group comprised between two and 11 positions (S6 Table). We then resampled genotype strings by choosing only one representative genomic position from each LD group, and did so for all possible combinations of positions. For example, if we found two LD groups comprising two and three genomic positions within a 250kbp window, respectively, we reconstructed 2×3=6 resampled nucleotide strings for this phenotype. This procedure resulted in between 2 and 288 different nucleotide strings for each phenotype, which we used to reconstruct genotype networks. We then calculated the frequency of epistasis and the fraction of accessible mutational paths for these genotype networks. Our observations are summarized in S6 Fig, which shows that epistasis is still pervasive, and that mutational paths still remain accessible. In summary, eliminating closely linked genomic position that may show LD from our analysis, does not affect our conclusions.

## S4 Text: Structure of the phylogenetic tree of all 199 A. thaliana accessions in Newick format

(((((6965:0.11947,8231:0.16595)1.000:0.07832,(6964:0.18878,(8242:0.14674,(7519:0.13946,7518:0.15036)1.000:0.05146)1.000:0.04362)1.000:0.03232)1.000:0.03791,(8230:0.23414,((6905:0.23630,(6921:0.00900,6920:0.00722)1.000:0.21734)1.000:0.07578,((9058:0.19397,(8351:0.18740,((6969:0.01934, 6968:0.01276)1.000:0.19509,(((6016:0.11244,6064:0.08910)1.000:0.03657,((6918:0.06291,6917:0.06659)1.000:0.07415,(6913:0.08372,6009:0.05839)1.000:0.06074)1.000:0.02762)1.000:0.04637,(8376:0.14489,((6043:0.01075,6046:0.01201)1.000:0.11112,(6900:0.01097,6901:0.00903)1.000:0.11060)1.000: 0.05878)1.000:0.02955)1.000:0.03646)1.000:0.03399)1.000:0.05312)1.000:0.03410,((7516:0.07581,75 17:0.06367)1.000:0.11888,6074:0.20963)1.000:0.04069)0.991:0.01976)1.000:0.02623)1.000:0.02508) 1.000:0.01468,(((8249:0.18284,8222:0.18690)1.000:0.04555,(6974:0.19817,(8241:0.16725,8326:0.186 29)0.981:0.03083)1.000:0.02005)1.000:0.02806,((6932:0.21420,7346:0.17669)1.000:0.04086,8283:0.2 2155)1.000:0.02786)0.702:0.01700)1.000:0.01398,(((8306:0.16102,(8423:0.14408,8426:0.15765)1.000:0.04764)1.000:0.05676,(8240:0.21050,(8237:0.16152,(8422:0.01677,8266:0.03585)1.000:0.13268)1.000:0.04415)1.000:0.03377)1.000:0.03048,((6946:0.21058,(8256:0.15720,(8258:0.01371,8259:0.01639)1.000:0.14409)1.000:0.07552)1.000:0.03083,(9057:0.20142,((8369:0.09961,(6040:0.06552,8335:0.05301)1.000:0.03632)1.000:0.06621,(6973:0.19563,8247:0.06266)1.000:0.07966)1.000:0.06442)1.000:0.04795)0.991:0.01567)1.000:0.01814,((((((6976:0.14105,6975:0.13967)1.000:0.06159,(8311:0.20332, 8314:0.24478)1.000:0.04596)1.000:0.02297,((((((7460:0.15684,(6984:0.13670,8285:0.14816)1.000:0.0 3621)0.000:0.01868,(8236:0.18716,8412:0.17107)1.000:0.04178)0.911:0.01573,((8284:0.13586,6008: 0.14782)1.000:0.04430,(6903:0.19826,(6985:0.14482,5837:0.15385)1.000:0.03647)0.709:0.01697)1.0 00:0.02149)1.000:0.02115,((7520:0.14893,7521:0.15200)1.000:0.05502,((8300:0.15512,7296:0.17518) 1.000:0.04614,(8378:0.18096,(8334:0.21031,8389:0.14127)0.954:0.03168)1.000:0.02091)0.949:0.01256)1.000:0.02139)1.000:0.01661,((8365:0.15800,7424:0.16439)1.000:0.04880,((6951:0.15981,(6956:0.07838,6957:0.06408)1.000:0.09396)1.000:0.03777,(8235:0.17347,8313:0.17866)1.000:0.04280)1.000: 0.01679)1.000:0.01604)0.299:0.01644,(7275:0.18838,6243:0.18354)1.000:0.04893)1.000:0.01914)1.0 00:0.02046,(((7163:0.23357,100000:0.16726)1.000:0.08911,((6978:0.16994,8325:0.18957)1.000:0.036 18,((8354:0.14909,(7438:0.14663,(7323:0.12776,((8424:0.09970,(6962:0.06085,(6963:0.07289,6929:0.04555)1.000:0.02373)1.000:0.03178)1.000:0.05498,(6931:0.10523,6930:0.10847)1.000:0.04124)0.99 9:0.02225)1.000:0.03316)1.000:0.05039)1.000:0.05134,6981:0.20754)1.000:0.02909)0.964:0.01902)1. 000:0.02494,(6980:0.19604,(6919:0.19334,(8388:0.13210,(8290:0.01137,6910:0.00990)1.000:0.11193)1.000:0.06397)1.000:0.03057)1.000:0.02748)1.000:0.01910)1.000:0.02168,(6042:0.22583,(7477:0.20 011,(6899:0.22983,(6972:0.01069,8395:0.00700)1.000:0.21031)0.975:0.04214)1.000:0.03121)1.000:0.02987)1.000:0.02518,(((((8343:0.20087,8270:0.15841)0.262:0.03910,6937:0.19158)1.000:0.02735,(6979:0.17373,7418:0.17832)1.000:0.04764)1.000:0.03675,((((8275:0.20273,(6977:0.19925,(6936:0.13298,6958:0.12347)1.000:0.07205)1.000:0.03337)1.000:0.04378,(((6959:0.13101,6960:0.12936)1.000:0.0 8989,(8329:0.17753,8213:0.18729)1.000:0.04352)1.000:0.02234,((((6924:0.15357,(6943:0.14815,((6966:0.08944,8254:0.05956)1.000:0.05280,(6944:0.09149,(7064:0.11884,6923:0.08907)1.000:0.02712)0.923:0.02009)1.000:0.05672)1.000:0.03757)1.000:0.04921,(8214:0.16623,(6709:0.02632,6908:0.00935)1.000:0.14562)1.000:0.03796)1.000:0.02758,((8215:0.16399,7306:0.19057)1.000:0.04927,(8245:0.01434,6926:0.01360)1.000:0.17812)1.000:0.02236)1.000:0.02665,8297:0.22455)1.000:0.02116)1.000:0.02542)1.000:0.01551,((6928:0.18188,(7523:0.18381,(6983:0.12115,((((8233:0.01042,7033:0.01003)1.000:0.00207,(6927:0.00884,7515:0.00838)1.000:0.00192)0.072:0.00149,7526:0.00729)1.000:0.03018, (7524:0.01278,7525:0.18466)1.000:0.05293)1.000:0.04924)1.000:0.04552)1.000:0.08863)1.000:0.06388,((6967:0.16390,6907:0.16951)1.000:0.05746,(6914:0.20681,7514:0.23933)1.000:0.04160)1.000:0.02656)1.000:0.02958)1.000:0.01785,(((8337:0.16086,(6904:0.17825,8243:0.14403)1.000:0.03995)1.000:0.05314,(6897:0.21090,(6906:0.18997,8353:0.16062)1.000:0.06259)0.174:0.02061)1.000:0.02107,((8264:0.18676,((6961:0.12867,8357:0.12810)1.000:0.04161,(6971:0.14387,(6970:0.11647,6933:0.11548)1.000:0.05097)1.000:0.02607)1.000:0.03050)1.000:0.06702,((7522:0.20472,(8274:0.16016,6911:0.19308)0.489:0.05560)1.000:0.06061,(6988:0.18099,7081:0.16653)1.000:0.05313)1.000:0.02916)1.000: 0.03863)1.000:0.01676)1.000:0.01813)1.000:0.02166,(((((7062:0.16342,7014:0.15879)1.000:0.07102,((8387:0.06028,6915:0.04404)1.000:0.14946,6982:0.20061)1.000:0.02664)1.000:0.02885,((8374:0.20226,7231:0.19056)0.812:0.03727,8239:0.25334)1.000:0.02703)1.000:0.02206,(((8312:0.21008,(8420:0.18422,7094:0.17982)1.000:0.04050)1.000:0.03499,((6940:0.17601,6942:0.19329)1.000:0.04298,((8411:0.00378,(8366:0.00798,6916:0.01542)0.568:0.00310)1.000:0.23309,(7000:0.20400,8271:0.19677)1.000:0.04001)1.000:0.02969)1.000:0.02426)1.000:0.01613,((8323:0.16721,6898:0.20127)1.000:0.07282,((7255:0.21219,(8296:0.22045,6939:0.17197)1.000:0.03889)1.000:0.01822,((6922:0.16277,(6909:0.12794,7461:0.09343)1.000:0.17047)1.000:0.04662,(7282:0.17849,7123:0.18348)1.000:0.03170)1.000: 0.02452)1.000:0.03398)1.000:0.02199)1.000:0.01900)1.000:0.02303,((8310:0.19672,8265:0.20580)1.000:0.04651,(7147:0.21169,(6945:0.17630,8430:0.18666)1.000:0.05292)1.000:0.02744)1.000:0.03546) 1.000:0.01914)1.000:0.02393)1.000:0.01878);

---

**S1 Fig.**
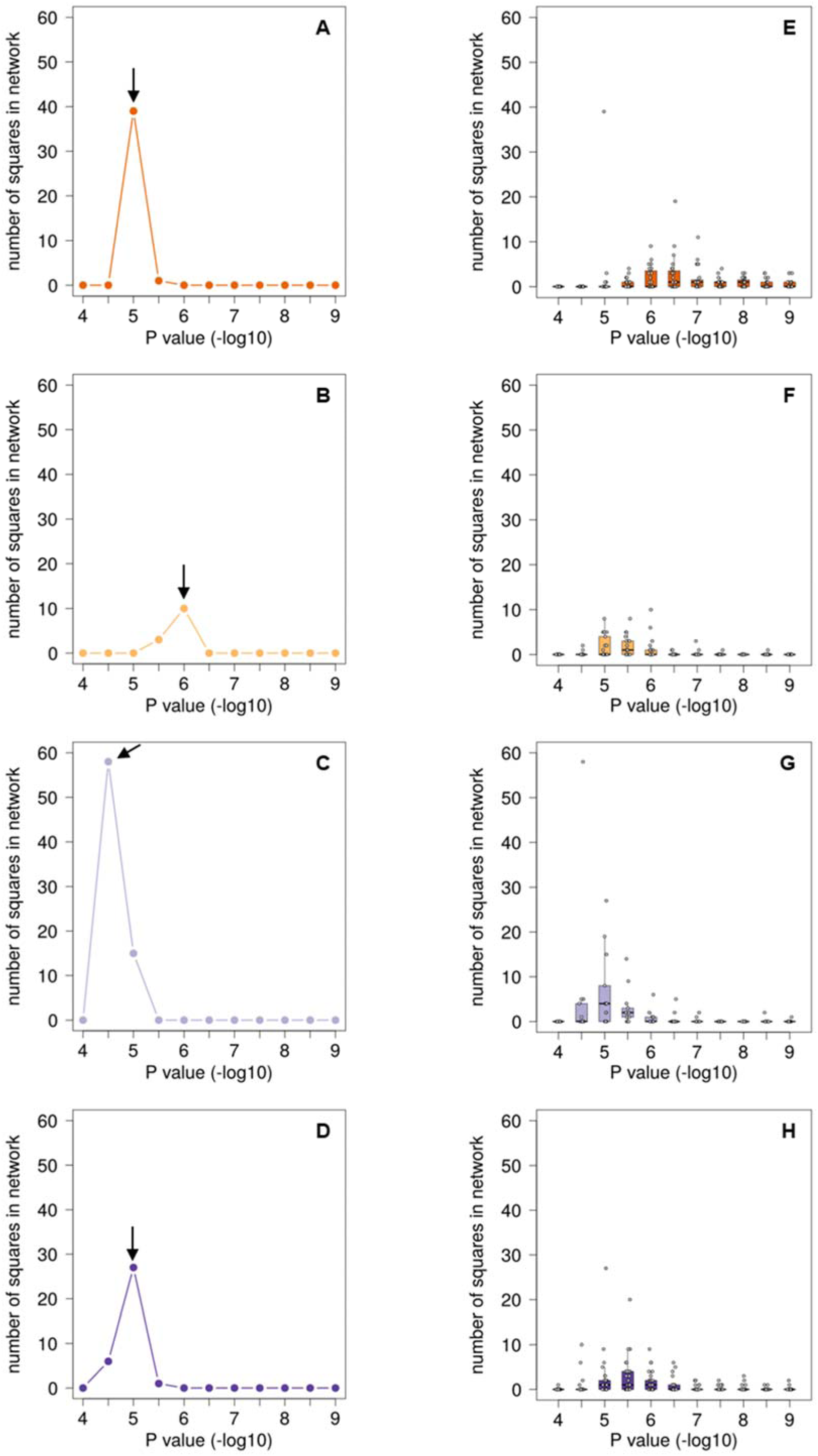
Relationship between *P*-value cutoff and number of squares in a genotype network. We asked which *P*-value cutoff between 10^-4^ and 10^-9^ (horizontal axes) results in the largest number of squares (vertical axes) for each of our 80 phenotypes. (A *–* D). For each of the four example phenotypes discussed in the main text, one single *P-*value cutoff leads to the largest number of squares. This *P*-value cutoff was 10^-5^ for the phenotype leaf diameter at flowering (A), 10^-6^ for the phenotype arsenic concentration (B), 10^-4.5^ for the phenotype bacterial growth (C), and 10^-5^ for the phenotype plant width (D). Arrows in each panel indicate the *P*-value that yields the largest number of squares in each of the four phenotype categories. (E *–* H) Box plots show the distribution of square number for all phenotypes in the categories flowering (E), ions (F), defense (G), and development (H). In each box plot, the central horizontal line indicates the median value, the lower box limit shows the value of the first quartile, the upper box limit depicts the third quartile, the whiskers indicate values within the 1.5-fold interquartile range, and each open circle represents one data point.

**S2 Fig.**
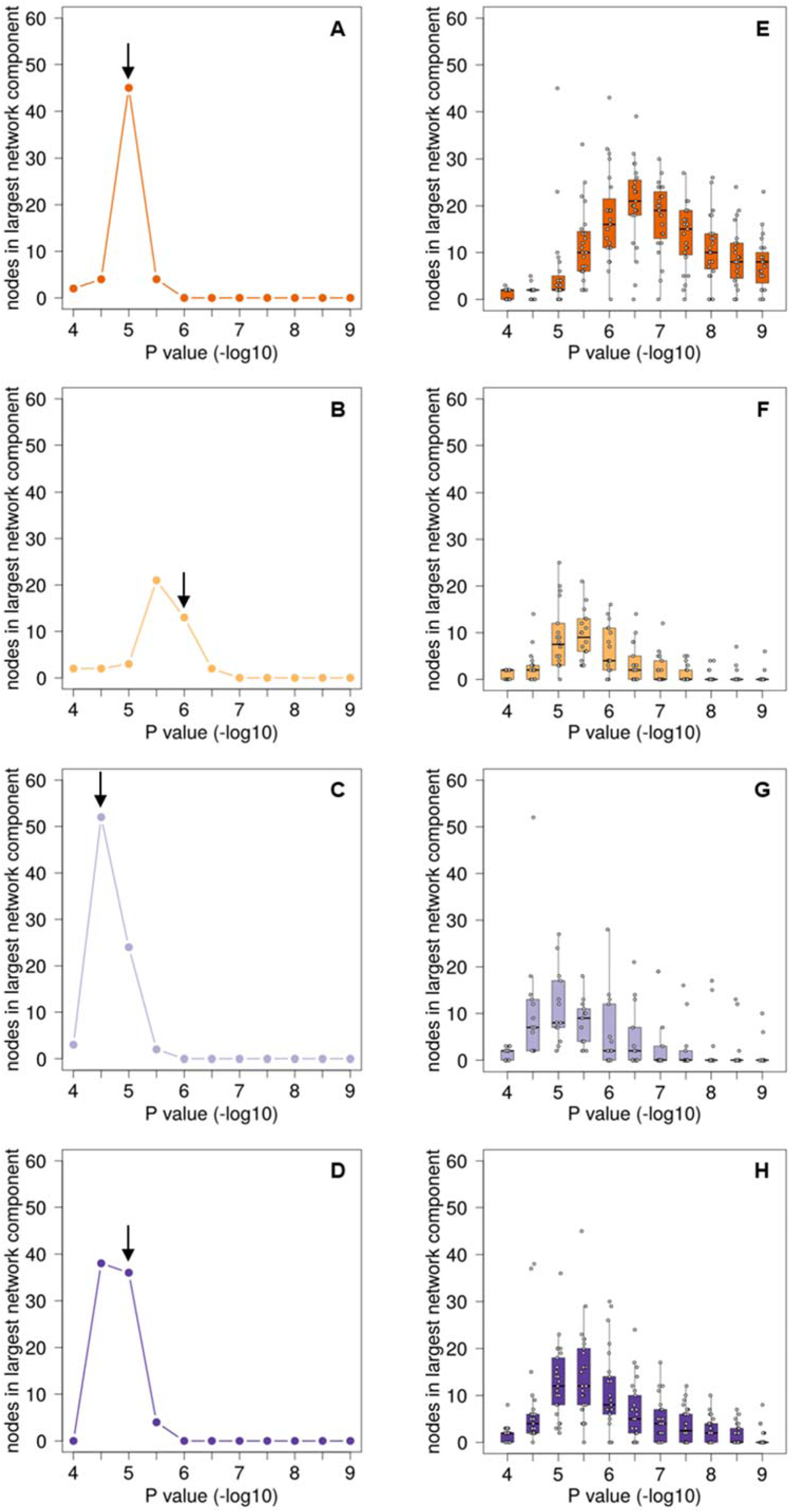
Relationship between *P*-value cutoff and size of largest genotype network component. We asked which *P*-value cutoff between 10^-4^ and 10^-9^ (horizontal axes) results in the largest connected component (vertical axes), and did so for each of our 80 phenotypes. Panels A to D show the results of the four example phenotypes discussed in the main text, namely leaf diameter at flowering (A), arsenic concentration (B) bacterial growth (C), and plant width (D). Arrows in each panel indicate the *P*-value that was chosen based on the number of squares in the genotype network, as shown in Fig S1. Panels E to H show box plots with the distribution of the sizes of largest network components for all phenotypes in the categories flowering (E), ions (F), defense (G), and development (H). In each box plot, the central horizontal line indicates the median value, the lower box limit shows the value of the first quartile, the upper box limit depicts the third quartile, the whiskers indicate values within the 1.5-fold interquartile range, and each open circle shows one data point.

**S3 Fig.**
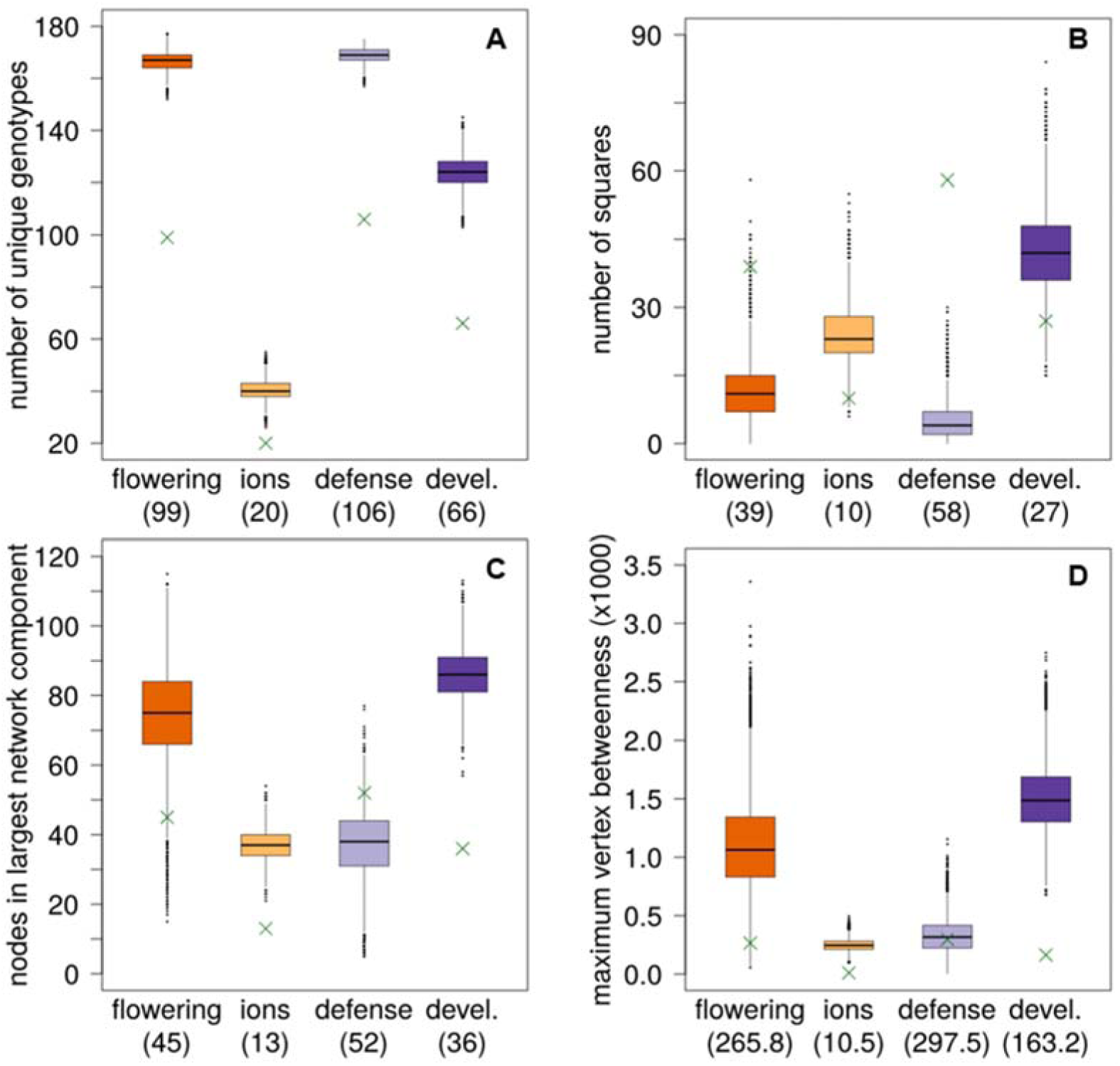
Properties of random genotype networks. We computed the number of unique genotypes (A), the number of squares in a genotype network (B), the number of vertices in the largest component with a size of at least two (C), and the maximum value of vertex betweenness (D) resulting from 10,000 randomly generated genotype networks. Box plots illustrate results from 10,000 replicates, and do so separately for each phenotype category (“devel”: development). Green crosses and numbers in parentheses on the horizontal axis indicate the empirically observed values for the example phenotypes in the flowering (plant diameter at flowering), ions (arsenic concentration), defense (bacterial growth) and development (“devel”, plant width) categories.

**S4 Fig.**
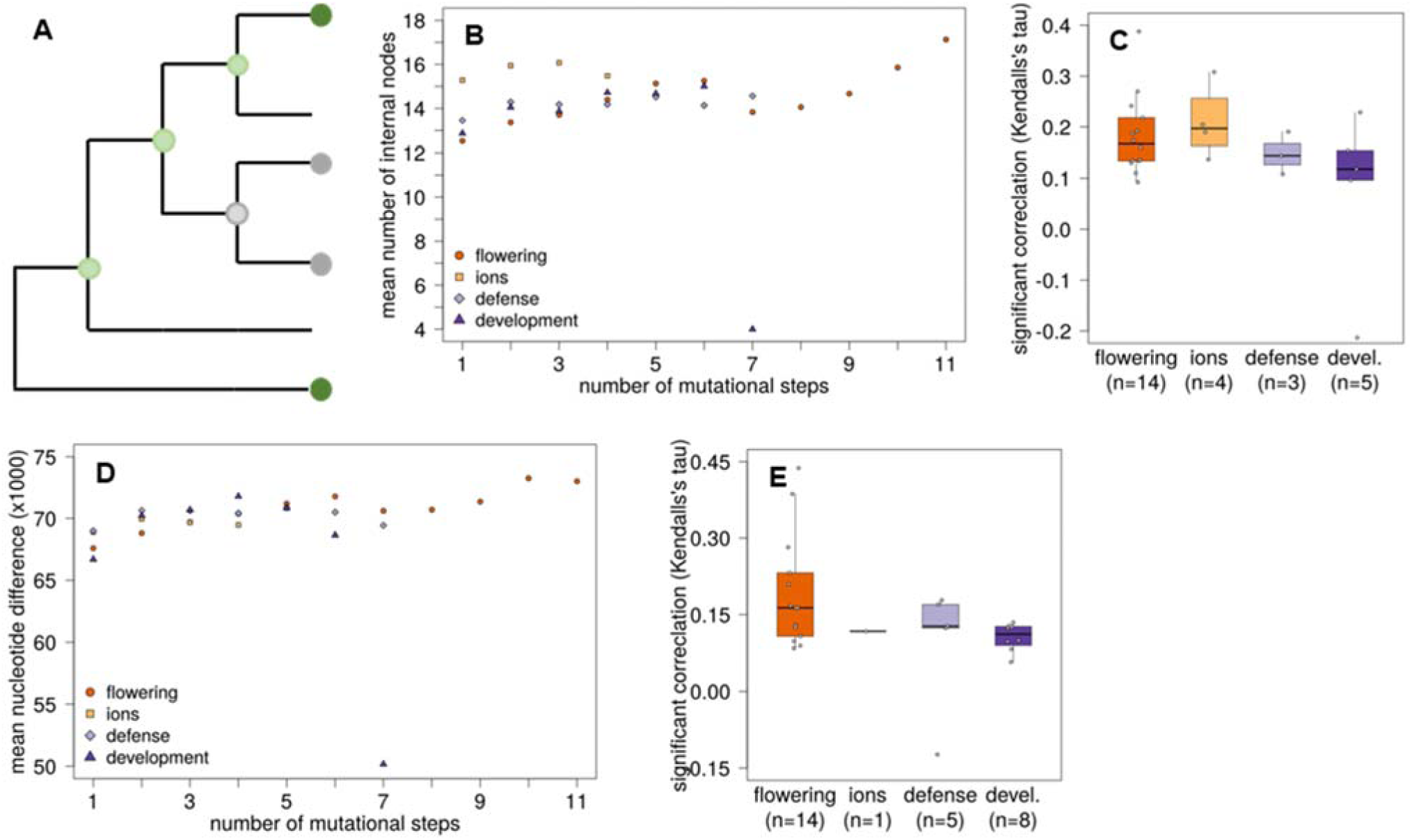
Relationship between mutational path lengths and phylogenetic distance. We compared the mutational path length between two genotypes (accessions) in our genotype networks with the number of branch-points separating these accessions on a phylogenetic tree. (A) Principle of counting internal branch-points. An accession (vertex) in the genotype network corresponds to one leaf on the phylogeny tree shown in S7 Fig. For example, the two leaves shown here as dark grey circles are separated by one internal branch-point (light grey circle), and the two leaves shown as dark green circles are three internal branch-points apart (light green circles). (B) For our four example genotype networks shown in Fig 2, we estimated the length of the shortest path between all vertices. We then sorted all path according to their length (horizontal axis) and calculated the mean number of internal branch-points between two accessions separated by a given path length (vertical axis). We did this individually for the phenotype plant diameter at flowering (dark orange circles), zinc concentration (light orange squares), bacterial growth (light purple diamonds), and plant width (dark purple triangles). None of the four phenotypes showed a strong association between the number of internal branch points and the length of a mutational path (Kendall’s tau between −0.021 and 0.388; *P*-values ≤ 0.0473). (C) We extended the same analysis to all 80 phenotypes. Each box plot summarizes rank correlation coefficients (Kendall’s tau) for one phenotype category (devl, development). In each box plot, the central horizontal line indicates the median value, lower and upper box limits show the first and third quartile, whiskers indicate values within the 1.5-fold interquartile range, and each open circle shows one data point. The number *n* of phenotypes in each category with correlations significantly different from zero is shown on the horizontal axes. Panels (D) and (E) are analogous to panels (B) and (C), but show the number of nucleotide differences between two accessions as a phylogenetic distance measure. Again, we observed mainly weak correlations (Kendall’s tau between −0.123 and 0.438; *P*-values ≤ 0.0418) between the mutational path length and the number of nucleotide differences.

**S5 Fig.**
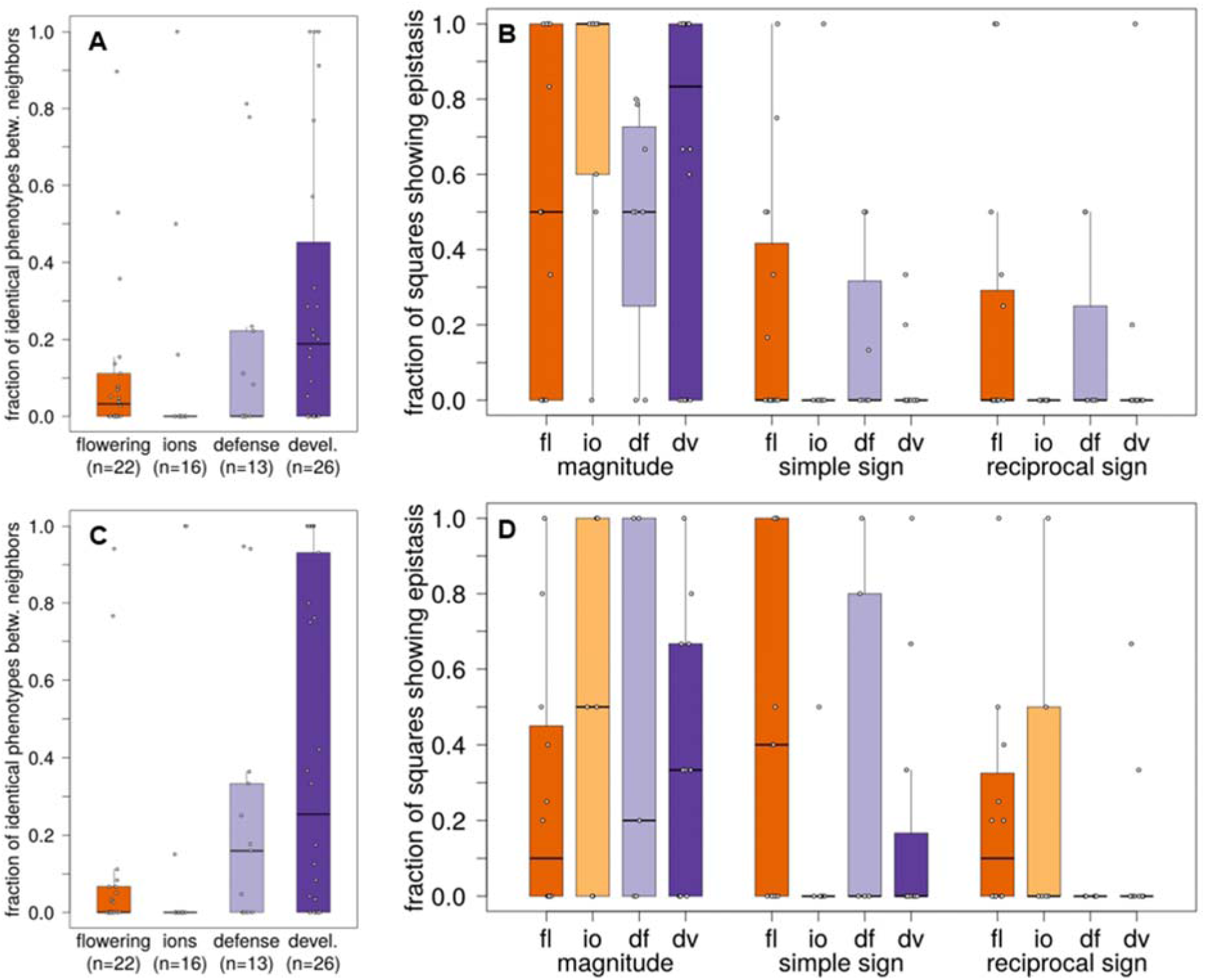
Fraction of identical phenotypes and prevalence of epistasis obtained with non-optimal phenotype *P*-value cutoffs. To determine how sensitive our quantification of epistasis is to the chosen *P*-value cutoff, we created genotype networks with non-optimal *P*-values for all phenotypes (*i.e*. *P*-values that do not yield the largest number of squares in a genotype network). Specifically, we used *P-*values whose -log_10_ value is higher (panels A and B) or lower (panels C and D) by 0.5 compared to the individual optimal *P*-value of each phenotype. (A and C) Fraction of genotype network neighbors with identical phenotypic values as a result of higher (panel A) or lower (panel C) *P*-values. We consider the absolute difference between the phenotypic values of two neighbors as identical if it does not exceed the coefficient of phenotypic variation (δ) for accessions with the same genotype at all phenotype-associated loci (Methods). We plot the fraction of neighbors with identical phenotypic values (vertical axis) in the form of a box plot for each of the four phenotype categories (“devl.”: development). The number of phenotypes in each category is shown on the horizontal axis as sample size (n). (B and D) Frequency of different types of epistasis for higher (panel B) or lower (panel D) *P*-values to reconstruct genotype networks. The vertical axis shows the fraction of squares with magnitude, simple sign, and reciprocal sign epistasis as a box plot, where data are grouped according to the type of epistasis and phenotype category (horizontal axis; “fl”: flowering; “io”: ions; “df”: defense; “dv”: development). In each box plot, the central horizontal line indicates the median value, lower and upper box limit shows the values of the first and third quartile, whiskers indicate values within the 1.5-fold interquartile range, and each open circle shows one data point.

**S6 Fig.**
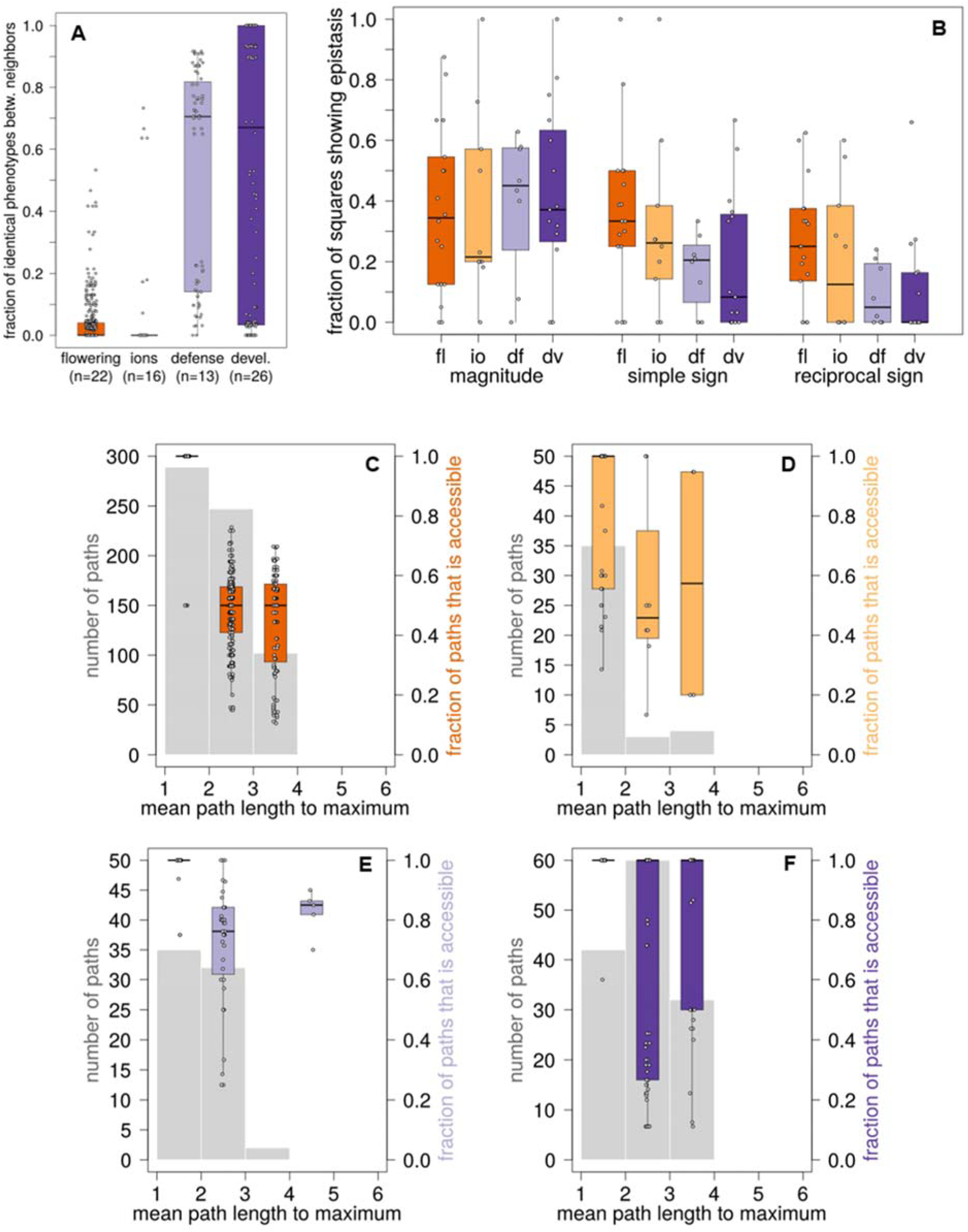
Incidence of epistasis and fraction of accessible mutational paths with correction for potential linkage disequilibrium. (A) Fraction of genotype network neighbors with identical phenotypic values. We plot the fraction of neighbors with identical phenotypic values (vertical axis) in the form of a box plot for each of the four phenotype categories (“devl.”: development). The number of phenotypes in each category is shown on the horizontal axis as sample size (n). (B) Frequency of different types of epistasis that we obtained when correcting for potential LD. The vertical axis shows the fraction of squares with magnitude, simple sign, and reciprocal sign epistasis as a box plot, where data are grouped according to the type of epistasis and phenotype category (horizontal axis; “fl”: flowering; “io”: ions; “df”: defense; “dv”: development). In each box plot, the central horizontal line indicates the median value, lower and upper box limit shows the values of the first and third quartile, whiskers indicate values within the 1.5-fold interquartile range, and each open circle shows one data point. (C – F) Distribution of mutational path lengths to maximum phenotypic values and fraction of accessible paths with correction for potential LD. For each phenotype, we calculated the mean length of the mutational path from each vertex in the network to the vertex with the maximum phenotypic value. We show the number of paths (left vertical axes) binned according to path length (horizontal axes, path length between 1 and 2 mutational steps, between 2 and 3 mutational steps, etc.) as indicated by the grey histograms. The colored box plot in each bin shows the fraction of paths to the maximum phenotypic value that is accessible (right vertical axes). Each panel shows the result for one phenotype category, namely flowering (C), ions (D), defense (E), and development (F).

**S7 Fig.**
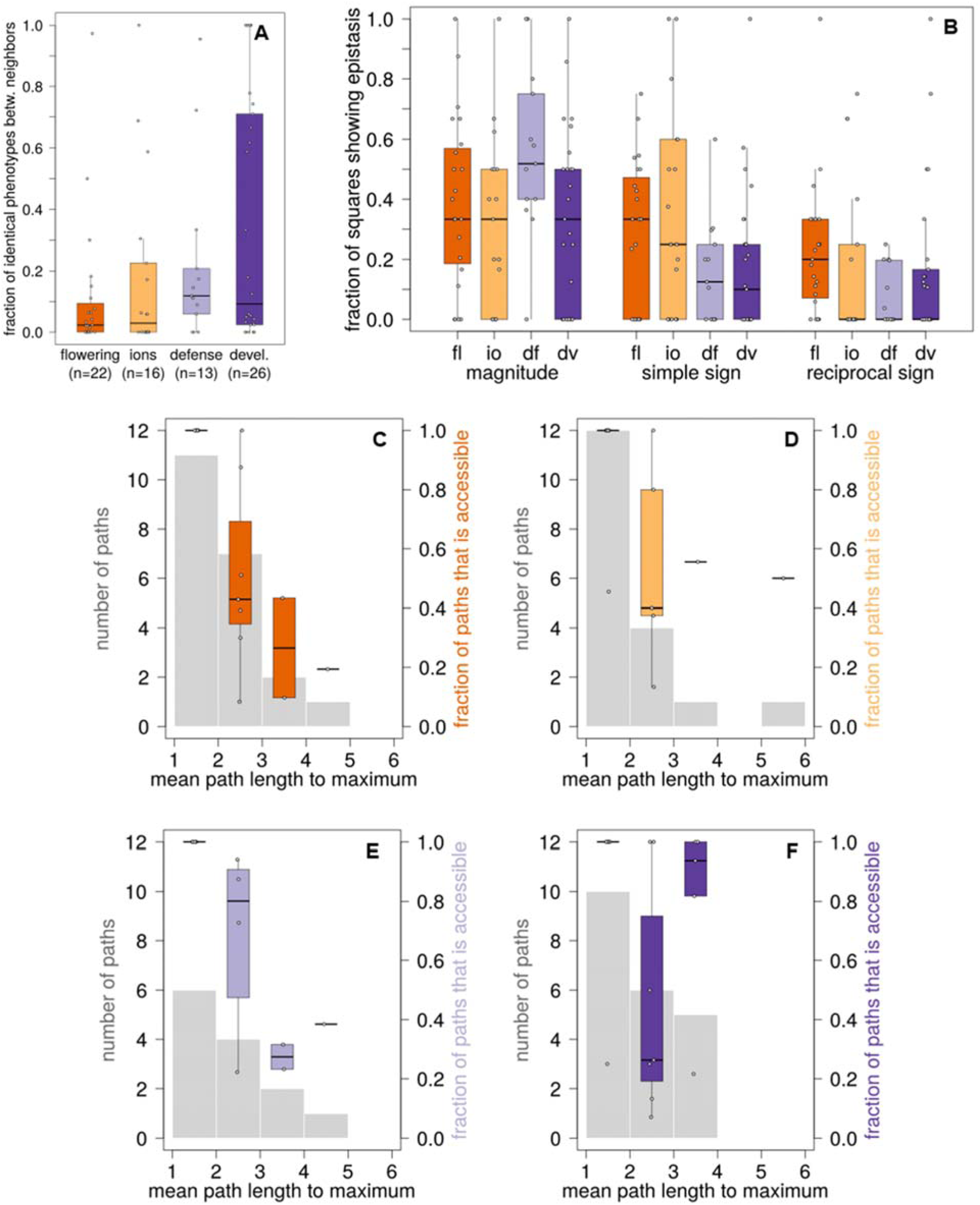
Frequency of epistasis and fraction of accessible mutational paths with correction for genomic positions with rare alleles (allele frequency below 5 percent). To determine how sensitive our findings of epistasis are to the presence of genomic positions with such alleles, we excluded all loci with such minor alleles from our data set before constructing genotype strings and genotype networks. Data are presented as described for S6 Fig.

**S8 Fig.**
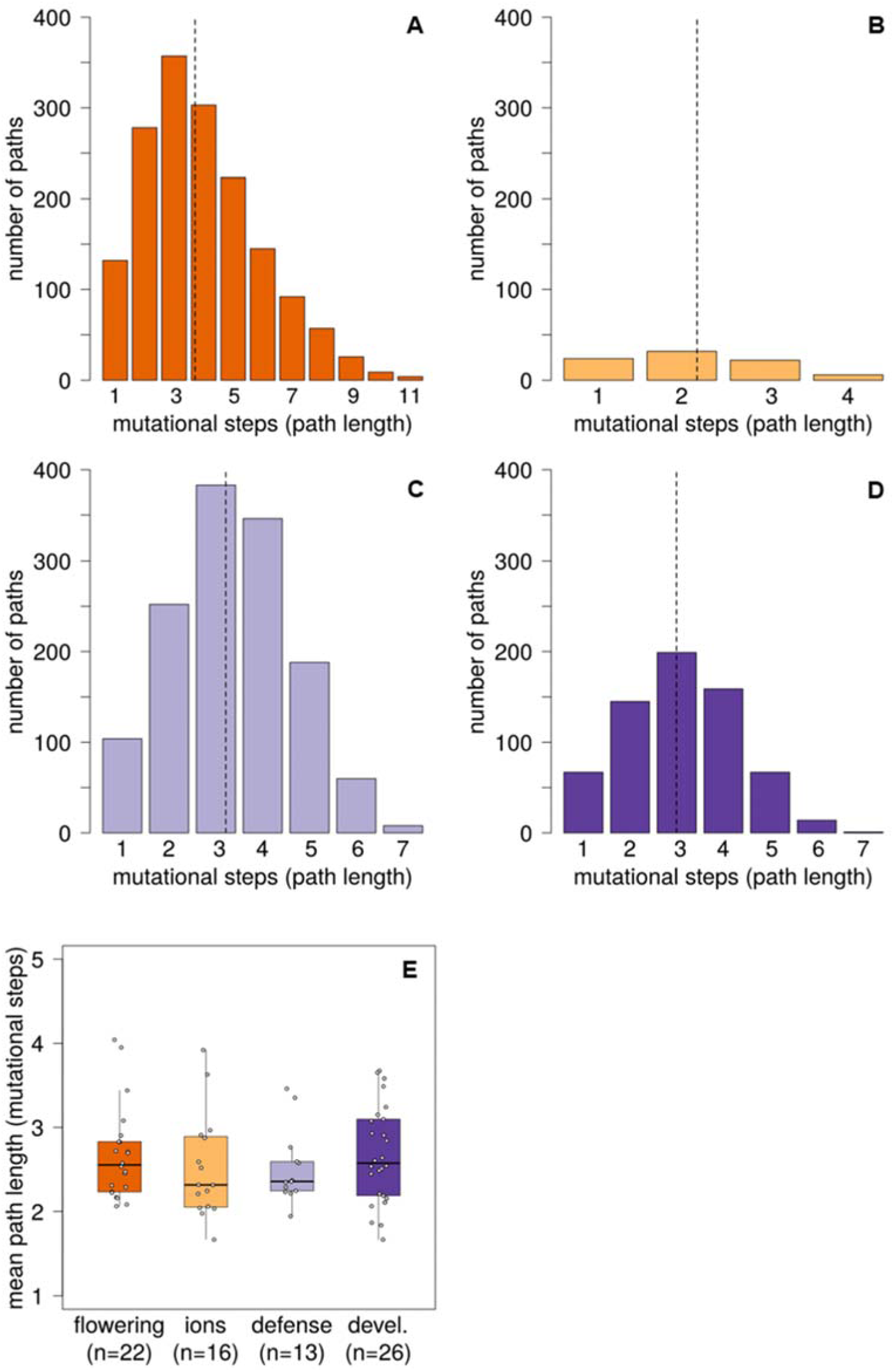
Distribution of all mutational path lengths. (A-D) Histograms show the distribution of mutational path lengths (horizontal axes) for all mutational paths present in the genotype networks of the four example phenotypes from the main text, i.e., for (A) plant diameter at flowering, (B) arsenic concentration, (C) bacterial growth and (D) plant width. Dashed vertical lines indicate the average mutational path length. (E) Mean mutational path lengths for all phenotypes in each of the four phenotype categories (horizontal axis, “devl.”: development). In each box plot, the central horizontal line indicates the median value, lower and upper box limits show the first and third quartile, whiskers indicate values within the 1.5-fold interquartile range, and each open circle shows one data point. The number of phenotypes in each category is shown as sample size (n).

**S9 Fig.**
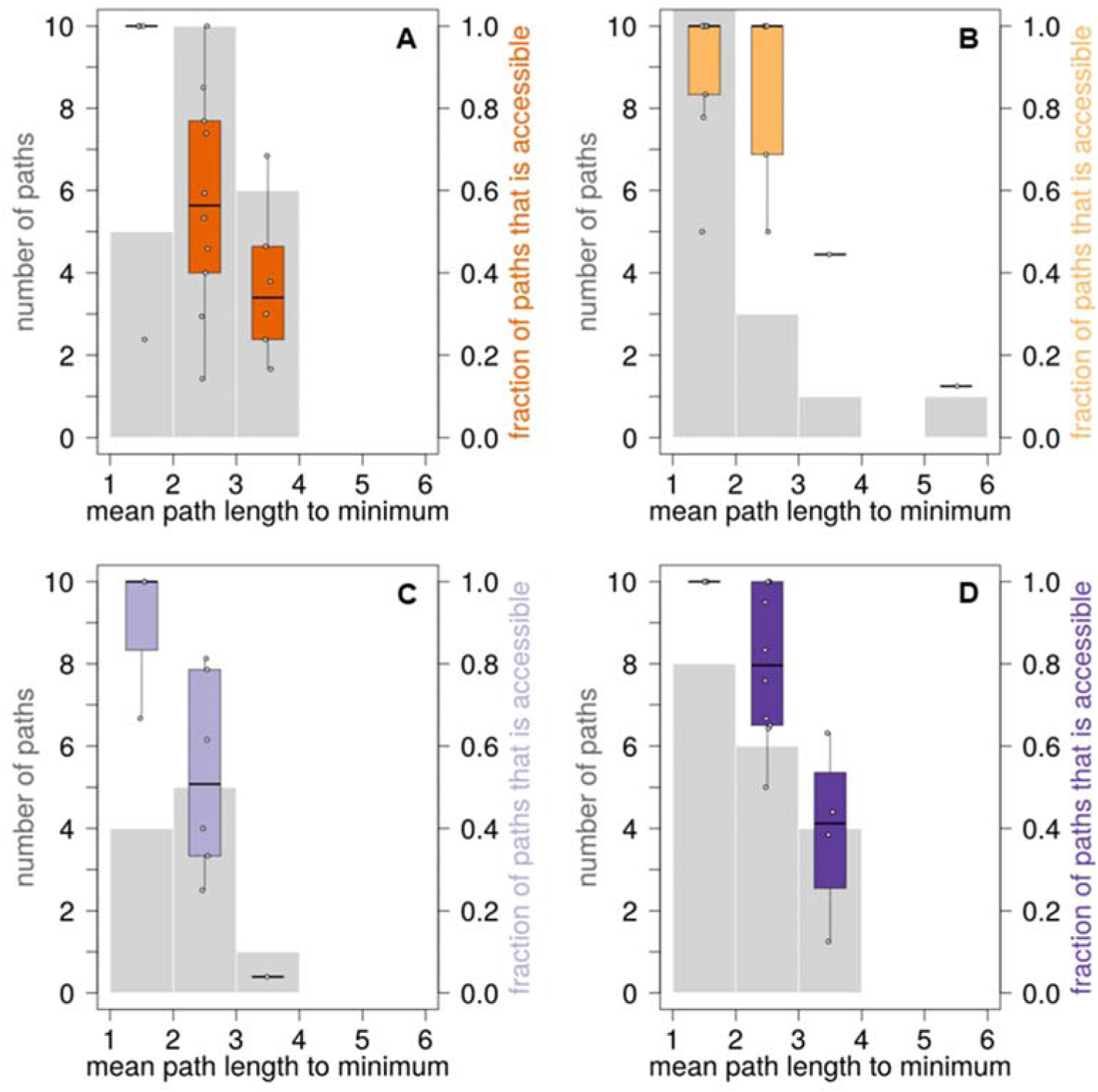
Distribution of mutational path lengths to minimum phenotypic values and fraction of accessible paths. For each phenotype, we calculated the mean length of the mutational path from each vertex in the network to the vertex with the minimum phenotypic value. We show the number of paths (left vertical axes) binned according to path length (horizontal axes, path length between 1 and 2 mutational steps, between 2 and 3 mutational steps, etc.) as indicated by the grey histograms. The colored box plot in each bin shows the fraction of paths to the minimum phenotypic value that is accessible (right vertical axes). Each panel shows the result for one phenotype category, namely flowering (A), ions (B), defense (C), and development (D).

**S10 Fig.**
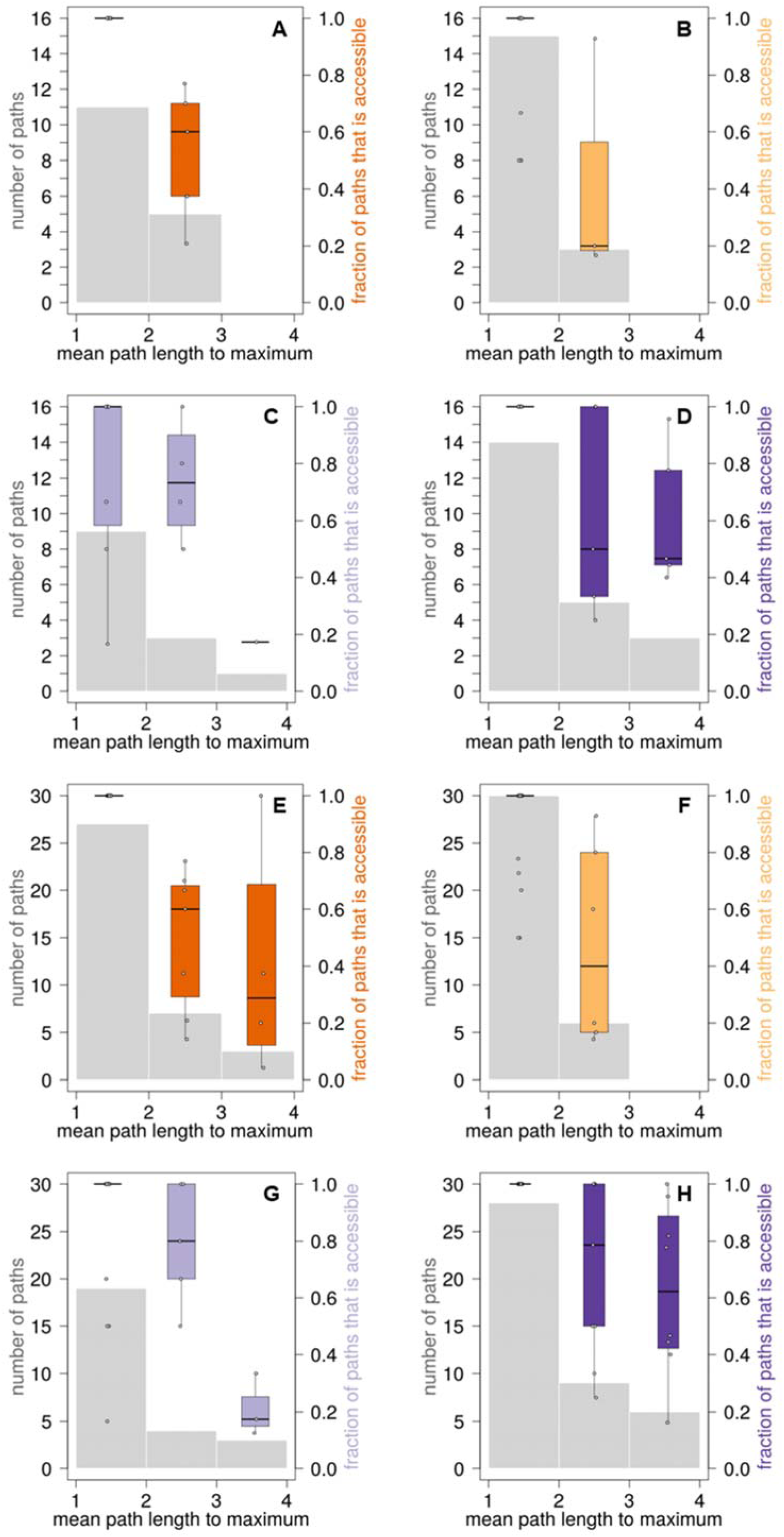
Distribution of mutational path lengths to maximum phenotypic values, and fraction of accessible paths in genotype networks that we created with non-optimal *P*-values (that is, *P*-values that do not yield the largest number of squares in a genotype network). For each phenotype, we reconstructed a genotype network based on *P*-values whose -log_10_ score is higher by 0.5 (panels A to D) or lower by 0.5 (panels E to H) than our default *P*-value for each phenotype. We then calculated the mean length of the mutational path from each vertex in all networks to the vertex with the maximum phenotypic value. We show the number of paths (left vertical axes) binned according to path length (horizontal axes, path length between 1 and 2 mutational steps, between 2 and 3 mutational steps, etc.) as indicated by the grey histograms. The colored box plot in each bin shows the fraction of paths to the maximum phenotypic value that is accessible (right vertical axes). Each panel shows the result for one phenotype category, namely flowering (A and E), ions (B and F), defense (C and G), and development (D and H).

**S11 Fig.**
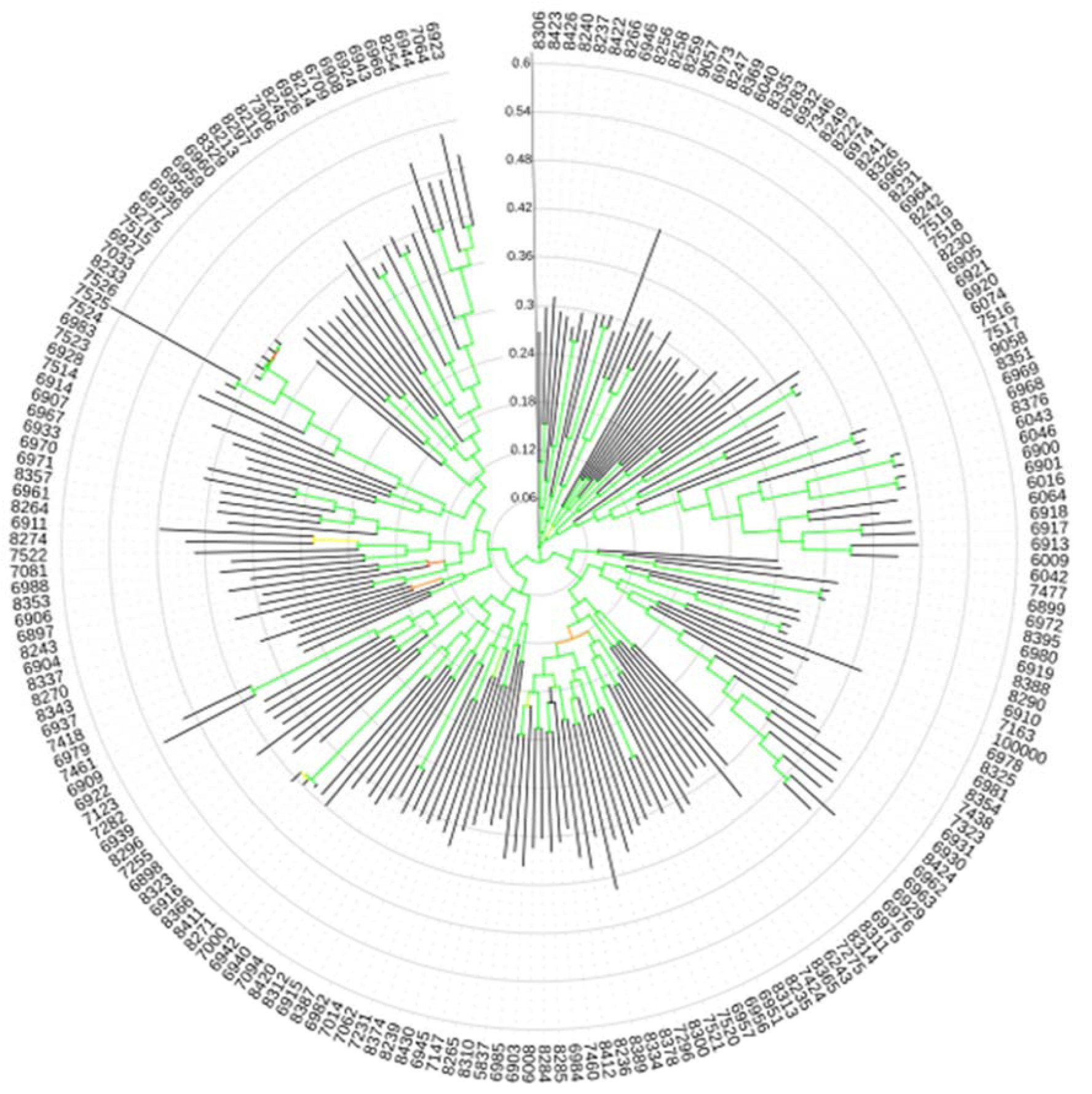
Phylogenetic tree of all 199 *A. thaliana* accessions considered in the present study. To construct this tree, we first concatenated nucleotides of polymorphic sites in the genomes that were represented on a re-sequencing chip [82]. All such strings had the same length and no position had missing information, that is, the concatenated strings can be aligned without gaps. We used the set of concatenated nucleotide strings as input to FastTree to create a phylogeny with 1,000 bootstraps, and displayed the resulting tree with iTOL. Each leaf corresponds to one accession and leaf labels indicate the accession number. Branches are colored according to their bootstrap support values. Green branches indicate a support value of 1, whereas yellow and orange branches indicate lower support values. Each inner grey circle indicates branch length in steps of 0.06 nucleotide substitutions per site, as indicated by the vertical scale shown in the upper part of the image.

